# A conserved opal termination codon optimizes a temperature-dependent tradeoff between protein production and processing in alphaviruses

**DOI:** 10.1101/2024.08.21.609082

**Authors:** Tamanash Bhattacharya, Eva M. Alleman, Alexander C. Noyola, Michael Emerman, Harmit S. Malik

## Abstract

Alphaviruses are enveloped, single-stranded, positive-sense RNA viruses that often require transmission between arthropod and vertebrate hosts for their sustained propagation. Most alphaviruses encode an opal (UGA) termination codon in nonstructural protein 3 (nsP3) upstream of the viral polymerase, nsP4. The selective constraints underlying the conservation of the opal codon are poorly understood. Using primate and mosquito cells, we explored the role and selective pressure on the nsP3 opal codon through extensive mutational analysis in the prototype alphavirus, Sindbis virus (SINV). We found that the opal codon is highly favored over all other codons in primate cells under native 37ºC growth conditions. However, this preference is diminished in mosquito and primate cells grown at a lower temperature. Thus, the primary determinant driving the selection of the opal stop codon is not host genetics but the passaging temperature. We show that the opal codon is preferred over amber and ochre termination codons because it results in the highest translational readthrough and polymerase production. However, substituting the opal codon with sense codons leads to excessive full-length polyprotein (P1234) production, which disrupts optimal nsP polyprotein processing, delays the switch from minus-strand to positive-strand RNA production, and significantly reduces SINV fitness at 37°C; this fitness defect is relieved at lower temperatures. A naturally occurring suppressor mutation unexpectedly compensates for a delayed transition from minus to genomic RNA production by also delaying the subsequent transition between genomic and sub-genomic RNA production. Our study reveals that the opal stop codon is the best solution for alphavirus replication at 37ºC, producing enough nsP4 protein to maximize replication without disrupting nsP processing and RNA replication transitions needed for optimal fitness. Our study uncovers the intricate strategy dual-host alphaviruses use at a single codon to optimize fitness.

## Introduction

Alphaviruses are a broad genus of enveloped, single-stranded, positive-sense RNA viruses that occur nearly worldwide and profoundly impact human health 1,2. Most extant alphaviruses are obligate dual-host viruses, *i*.*e*., their transmission relies on obligate alternation between arthropod vectors and vertebrate hosts 3. Dual-host alphaviruses are subject to divergent selective pressures in insect and vertebrate hosts, resulting in viral genomic features that might be host-specific ^4,5^. One such host-specific genome feature is the opal (UGA) termination codon that disrupts most alphaviruses’ non-structural polyprotein (nsP) open reading frame ^6^. The nsP3 opal codon is located at the C-terminal end of the non-structural protein 3 (*nsP3*) gene, just upstream of the non-structural protein 4 (*nsP4*) (Figure 1A) ^6^. First described in the Sindbis virus (SINV) genome, the opal codon is part of a Type-II programmed ribosomal readthrough (PRT) motif, which is characterized by the UGA codon followed by a cytosine (C), with a more modest preference for uracil (U) residues at the -2 and -3 positions (Figure 1A). These flanking nucleotides promote efficient PRT, during which cellular tRNAs outcompete canonical eukaryotic translation termination factors such as eRF1 and incorporate a sense codon instead of the termination codon ^7,8^. Through PRT, the *nsP* genes of most alphaviruses encode two distinct polyproteins ^9^. The first comprises nsP1-nsP2-nsP3 (P123). Ribosomal readthrough of the opal stop codon also produces lower amounts of the longer nsP1-nsP2-nsP3-nsP4 polyprotein (P1234) at a rate of approximately 5-20% ^10^. Since nsP4 is the viral RNA-dependent RNA polymerase (RdRp), sufficient PRT is required for *de novo* viral RNA synthesis.

**Figure 1.**
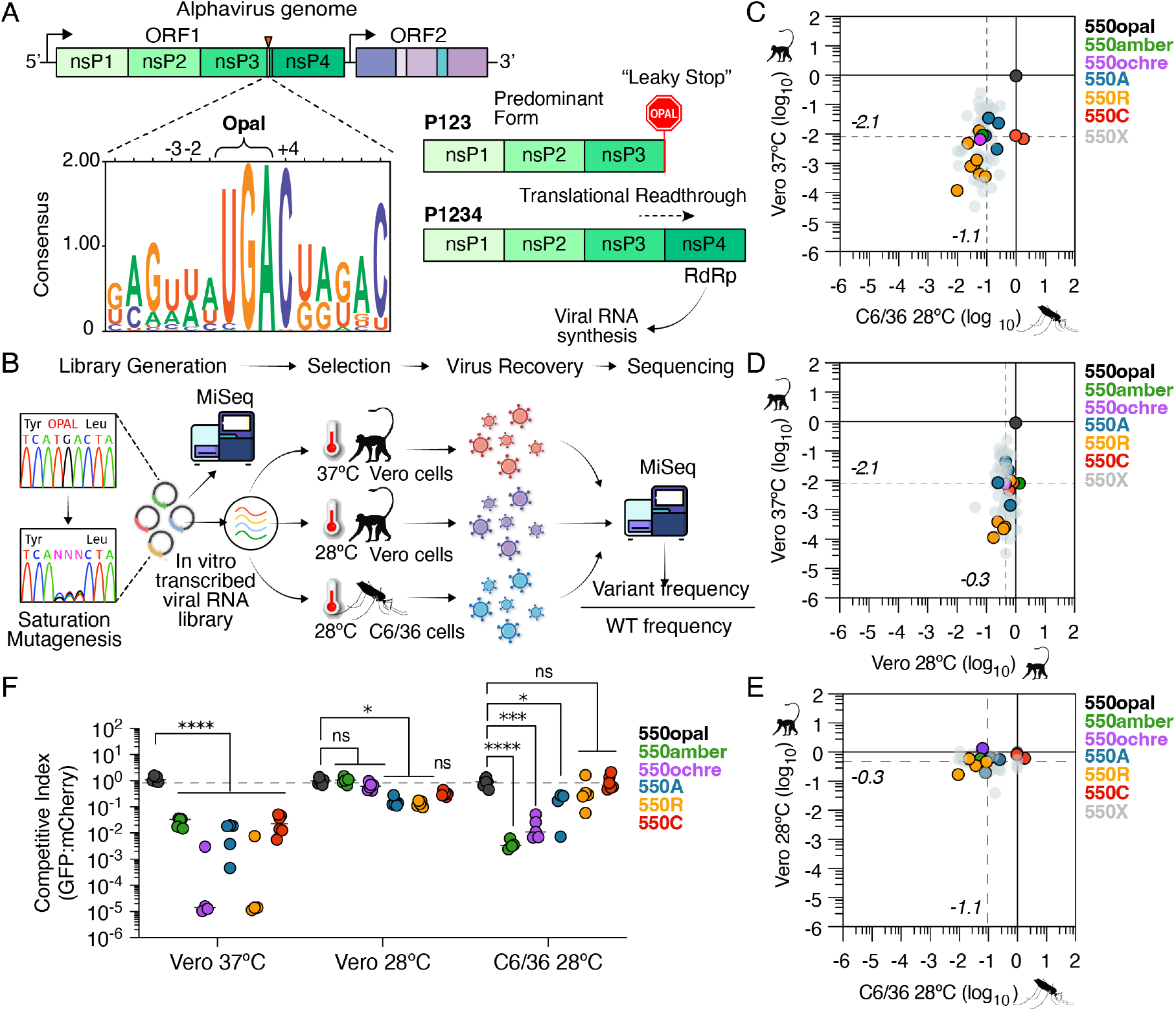
Saturation mutagenesis identifies host- and temperature-specific constraints acting on SINV nsP3 opal codon. (A) Schematic of alphavirus non-structural polyprotein indicating the location of the opal stop codon in nsP3. Logo plot shows conservation at the opal stop codon site and its genomic context among dual-host alphaviruses. -3 and +4 positions indicate constrained residues that aid in the translational readthrough of the opal codon. Translational readthrough of the “leaky” opal stop codon leads to nsP4 (RdRp) expression and commencement of *de novo* viral RNA synthesis. (B) Schematic of saturation mutagenesis screen. The WT opal stop codon in SINV nsP3 was mutated to one of 64 possible codons using PCR mutagenesis. The mutant virus library was transcribed *in vitro* and subjected to selection by passaging through African green monkey (Vero) cells grown at 37ºC or 28ºC and mosquito (C6/36) cells grown at 28ºC. Following selection, viruses were sequenced alongside the input/pre-selection plasmid library to determine the relative selection of each codon variant relative to the WT opal codon. (C-E) Correlation plot of mean ‘selection scores’ of codon variants across distinct environmental variables. Dotted horizontal and vertical lines depict the median selection scores for all non-opal codon variants. 550X refers to all sense codons excluding Alanine, Arginine and Cysteine codons. (F) Competition assays between GFP-tagged variant and mCherry-tagged wildtype (550opal) SINV strains. Two-way ANOVA with Tukey’s multiple comparisons test. ^****^ = P < 0.0001, ^***^ = P < 0.001, ^**^ = P < 0.01, ^*^ = P < 0.05, ns = not significant.

Since nsP processing and nsP4 expression are often rate-limiting steps for alphavirus replication, the almost strict preservation of the nsP3 opal codon suggests that the opal codon must have been conserved due to selective constraints. For example, previous analyses of SINV, Chikungunya virus (CHIKV), and O’nyong-nyong virus (ONNV) isolates with opal versus ‘sense’ (non-stop) codons revealed significant differences in tissue tropism (CHIKV, SINV), virulence (CHIKV, SINV), and vector competence (ONNV) ^11–13^. Replacing the opal codon with sense codons significantly reduced CHIKV replication kinetics, host-specificity, and virulence in mice ^12^. *In vitro* passaging experiments with Eastern Equine Encephalitis Virus (EEEV), CHIKV, and ONNV further demonstrated the selective consequences of opal-to-sense codon mutations ^14–16^. For instance, experimental passaging of the alphavirus EEEV exclusively in mosquito cells resulted in the opal codon being replaced by sense codons in multiple independent lineages ^14^. In contrast, opal-to-sense codon substitutions were not observed following EEEV passaging in vertebrate cells ^14^. Similarly, Semliki Forest virus (SFV) and CHIKV variants encoding either sense codons (CGA, arginine) or alternate stop codons (UAG, amber; UAA, ochre) were less fit in vertebrates but more tolerated in mosquito cells ^17,18^. These and related studies led to the hypothesis that host-specific constraints have preserved the opal stop codon in alphaviruses. Yet, the exact nature of these constraints and how they affect alphavirus fitness remains unknown.

Here, we investigated the function and selective constraint on the nsP3 opal codon using comprehensive mutational scanning of this codon in the prototype alphavirus Sindbis virus (SINV) in both primate (Vero) and mosquito (C6/36) cells. We find that the opal codon outperforms all other codon variants in Vero cells at 37ºC, but this strong preference is reduced in mosquito cells and Vero cells at 28ºC (passaging temperature for mosquito cells). Thus, the primary determinant selecting for the opal stop codon is not host genetics but viral passaging temperature. We demonstrate that the opal codon is favored over the amber and ochre stop codons because it provides the highest translational readthrough and production of the nsP4 protein at 37ºC. However, the replacement of the opal codon with sense codons produces too much full-length polyprotein (P1234), which impairs optimal nsP polyprotein processing, leading to a delay in the switch between minus-strand and positive-strand production, which drastically reduces SINV replicative fitness at 37ºC. A naturally occurring ‘suppressor’ mutation in a SINV strain carrying a sense codon unexpectedly compensates for a delayed transition from minus to genomic RNA production by further delaying the subsequent transition between genomic and sub-genomic RNA production.

Our studies reveal that the opal stop codon is the optimal solution to an inherent tradeoff in alphavirus replication at 37ºC. It produces enough nsP4 protein to maximize SINV replication without impairing nsP processing or synchronized RNA replication transitions — minus-strand to genomic to sub-genomic RNA synthesis — required for optimal fitness. Our findings reveal the mechanistic basis of an exquisite strategy deployed by dual-host alphaviruses at a single codon to optimize fitness in disparate hosts at their native temperatures.

## Results

### Host temperature imposes the predominant selection pressure for the nsP3 opal codon

To investigate the selective constraints acting on the in-frame opal codon site in alphavirus nsP3 (Figure 1A) in different hosts and at different temperatures, we generated a saturation mutagenesis library of all possible mutations at the nsP3 opal stop codon site (codon 550) of the prototype alphavirus Sindbis (SINV). A similar strategy was previously used to identify host-specific constraints in other alphavirus proteins ^19,20^. Using *in vitro* transcription, we created replication-competent viral RNA from this library and transfected it either into Vero cells (derived from African green monkeys) maintained at 37ºC (hereafter referred to as Vero-37º) or into C6/36 cells (derived from *Aedes albopictus* mosquitoes) maintained at 28ºC (hereafter referred to as C6/36) (Figure 1B). We deliberately chose vertebrate and mosquito cells with compromised innate immunity to remove the dominant effects of selective pressure imposed by vertebrate-specific Type-I/II IFN or mosquito-specific RNA-interference pathways ^21,22^. To further distinguish between the effects of temperature and host genetics, we also transfected the library into Vero cells at 28ºC (or Vero-28º) (Figure 1B); reduced cell viability precluded us from testing the variant library in C6/36 cells at 37ºC. Seventy-two hours post-transfection, we harvested viruses from cellular supernatants and conducted deep sequencing to determine ‘selection scores’ for each SINV variant. These selection scores represent the ratios of variant frequency post-selection to their initial frequencies in the input library normalized to the wildtype opal codon (Figure 1B, also see Materials and Methods). Thus, selection scores quantify the relative enrichment and depletion of SINV variants as indicators of their replicative fitness, with positive selection scores indicating enhanced fitness and negative selection scores indicating reduced fitness (Figure 1B).

Consistent with its evolutionary conservation, we found that the opal codon variant significantly outperformed all other codon variants in Vero-37º (Figure 1C, Suppl. Figure 1A-B), with a mean selection score for all non-opal variants of -2.1 ± 1.1 (on a log_10_ scale). Most sense codons and alternate termination codons – amber (UAG) and ochre (UAA) – are significantly depleted. Selection scores for synonymous codons encoding the same amino acid were similar, indicating that selection primarily acts at the protein level rather than the underlying viral RNA sequence (Suppl. Figure 1B). The opal codon remained the most preferred codon even in C6/36 cells, outperforming all codons except the cysteine sense codons (Figure 1C). However, the mean selection score of -1.1 ± 0.47 (on a log_10_ scale) for all non-opal codons is much less severe in C6/36 cells than the mean selection score in Vero-37º. These results show that the selective constraint on the SINV opal codon is less stringent in mosquito cells than in Vero-37º.

The dramatic difference in selective constraint acting on opal stop in Vero-37º versus C6/36 cells could result from genetic or metabolic differences between the host cell types or different temperatures of passaging. We, therefore, considered the possibility that differential selection scores reflect differences in tRNA abundance between host cell types. However, selection scores do not appear to correlate with codon usage frequency in host cells (*Chlorocebus aethiops* versus *Aedes albopictus*, Suppl. Figure 1C) ^23,24^. A previous study showed that SFV variants with arginine sense codons had lower fitness in chicken cells at 39ºC, but fitness could be restored at 30ºC in the same cells ^18^. To test for temperature-dependent effects more comprehensively, we compared selective constraints acting on this codon in Vero-37º versus Vero-28º (Figure 1D, Suppl. Figure 1A-B). Remarkably, we found that the mean selection score was only -0.34 ± 0.24 (on log_10_ scale) for all non-opal variants in Vero-28º; this represents the weakest preference for the opal codon among all three conditions tested. Several sense codons (including UGC for cysteine) and the amber stop codon (UAG) had identical selection scores to the opal stop codon in Vero-28º (Figure 1D, Suppl. Figure 1). As a result, there are only modest differences between C6/36 and Vero-28º cells (Figure 1E). These findings show that temperature, rather than host cell type differences, is the predominant factor driving selective retention of the SINV nsP3 opal codon.

We validated the selection scores obtained from our mutagenesis and pooled passaging strategy using two orthogonal assays. These assays measured either competitive or independent fitness of six SINV variants encoding either of three alternate stop codons — opal, amber, or ochre — or three sense-codons — arginine (550R), cysteine (550C), or alanine (550A). In the first validation assay, we measured the fitness of GFP-tagged SINV variants relative to a mCherry-tagged wildtype SINV strain (WT, with an opal stop) using co-infections in Vero-37º, Vero-28º, or C6/36 cells. We harvested cells 48 hours post-infection and analyzed them using flow cytometry to calculate the ratio of cells infected with GFP-tagged or mCherry-tagged SINV. We normalized the competitive index by comparing the ratio of GFP: mCherry cells in a control co-infection with GFP-tagged WT SINV (Figure 1F). We then quantified the competitive fitness of the other five GFP-tagged SINV variants relative to the mCherry-tagged WT SINV. We found that the 550opal variant outcompetes 550amber and 550ochre by two to four orders of magnitude in Vero-37º and C6/36 cells, but the relative fitness of 550amber and 550ochre is almost fully restored in Vero-28º. Similarly, the high fitness advantage of the 550opal variant over 550R, 550C, or 550A in Vero-37º is considerably reduced in C6/36 cells or Vero-28º, with 550C nearly identical in fitness to WT SINV in C6/36 cells.

To assess the independent fitness of the SINV variants, we also compared the infection rates of all SINV variants relative to wildtype SINV (550opal) in Vero-37º, Vero-28º, and C6/36 cells using flow cytometry (Suppl. Figure 2). The results from this assay were also consistent with those obtained from the pooled and pairwise competition assays. Thus, the selection scores we obtained from our pooled screening of all variants at *nsP3* codon 550 strongly correlate with individually measured competitive or absolute fitness measurements of the six selected SINV variants (Pearson R = 0.915, 95% CI = 0.7824 to 0.9683, P-value < 0.0001). These results confirm that the pooled selection scores are reliable measures of viral fitness, and that the WT SINV (550opal) significantly outperforms alternate stop and sense-codon variants in Vero-37º.

### Translational readthrough efficiency underlies fitness differences between alternate stop codons

We investigated why the opal stop codon is preferred over other stop codons in nsP3. All SINV variants encoding stop codons require translational readthrough to produce the nsP4 RdRp (RNA-dependent RNA polymerase), which is absolutely required for viral replication. Different stop codons have variable translational readthrough rates in eukaryotic mRNAs, with the opal codon exhibiting the highest readthrough frequency, followed by amber and then ochre ^25,26^. Since alphaviruses utilize host translation machinery, we reasoned that viral RNAs must be subject to similar translational preferences.

We designed dual-luciferase reporter constructs to measure the amount of readthrough with different stop codons on nsP3 (Figure 2A). The first reporter is Nano luciferase (Nluc) fused to *nsP3* upstream of codon 550 site. The second reporter is the Firefly luciferase (Fluc) reporter fused to *nsP4* downstream of the termination codon readthrough element, which is located in the first 100bp of nsP4 and is necessary for optimal readthrough (Suppl. Figure 3A) ^8,27,28,^. We transfected SINV P34 constructs driven by the Cytomegalovirus (CMV) promoter-enhancer in Vero-37º or Vero-28º and quantified Nluc and Fluc expression after 48 hours. We used analogous constructs driven by the Drosophila Actin (Act5C) promoter in C6/36 cells. We measured Fluc: Nluc ratios for all three stop codons in the SINV P34 construct to quantify translational readthrough efficiency, comparing Fluc: Nluc levels for a sense codon (alanine, GCA) that requires no translational readthrough. We found that the P34 construct encoding 550opal had ∼15% translational readthrough (relative to the sense codon) in Vero-37º (Figure 2B); these levels accord well with previous measurements of 5-20%^10^. In contrast, translational readthrough for the P34-550amber (5%) and P34-550ochre constructs (2%) was significantly lower in Vero-37º. Lower nsP4 expression was also apparent in western blot analyses of infected Vero-37º cells (Suppl. Figure 3C). These results clearly demonstrate a hierarchy in translational readthrough efficiency of the three termination codons within the context of the alphavirus genomic RNA.

**Figure 2.**
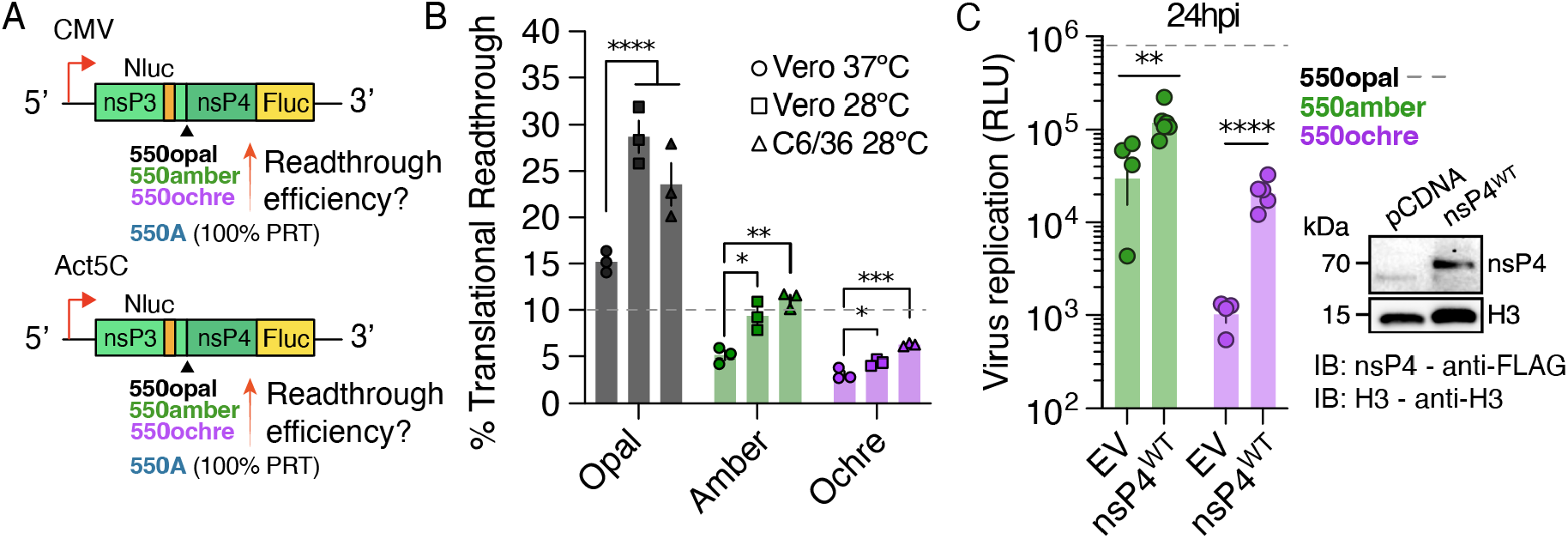
Temperature-dependent translation readthrough efficiency dictates the fitness of SINV variants encoding alternate stop codons. (A) SINV nsP3/4 translational read-through was quantified using dual-luciferase reporter constructs carrying CMV promoters in vertebrate cells and Act5 promoters in insect cells. (B) Translational readthrough of SINV variants containing alternate stop codons in Vero-37º, Vero-28º, or C6/36 cells. The horizontal dotted line denotes a 10% translational readthrough. (C) SINV variants carrying alternate stop codons were used to infect Vero-37º cells in the presence or absence (EV-empty vector) of wildtype SINV nsP4 *in trans*. Each virus expresses an in-frame Nluc reporter, whose expression is directly proportional to virus replication. Replication of SINV carrying alternate stop codons was quantified in the presence and absence of exogenous nsP4 using Nluc expression as a readout. Dotted line represents level of wildtype (550opal) SINV replication. Two-way ANOVA with Tukey’s multiple comparisons test. ^****^ = P < 0.0001, ^***^ = P < 0.001 ^**^ = P < 0.01, ^*^ = P < 0.05.

We found that translational readthrough efficiency for all stop codon variants was higher in both Vero-28º and C6/36 cells, although the hierarchy in translational readthrough efficiency (opal> amber> ochre) remained true in all conditions (Figure 2B). These results confirm that the disparities in replicative fitness observed between the wildtype (550opal) and the alternate stop codon variants 550amber and 550ochre) can be attributed to lower translational readthrough frequencies, which lead to reduced nsP4 protein expression. This discrepancy is particularly pronounced at 37ºC but is ameliorated at 28ºC, where higher readthrough rates for the amber and ochre codons partially rescue fitness defects. Given that infection rates of 550amber are at wildtype levels in Vero-28º cells (Suppl. Figure 2), we further conclude that a threshold level of ∼10% readthrough efficiency is sufficient to restore SINV fitness in vertebrate cells (Figure 2B). However, a higher readthrough frequency might be required to achieve wildtype levels of SINV fitness in mosquito cells, given the relative instability of nsP4^29^.

If replication defects of alternate stop codon SINV variants in Vero-37º were primarily due to insufficient nsP4 expression, we hypothesized that additional, exogenous nsP4 production might improve replicative fitness of 550amber and 550ochre SINV variants. We tested this hypothesis by co-transfecting a wildtype nsP4-expressing construct and the SINV variants. We found that exogenous nsP4 expression significantly enhanced the replication of both 550amber and 550ochre SINV variants (Figure 2C). Our findings demonstrate that lower translational readthrough and nsP4 production are the predominant cause of lower fitness of alternate stop codons at 37ºC, which is naturally ameliorated by higher translational readthrough at 28ºC.

### Fitness loss of sense-codon SINV variants is due to delayed plus-strand (genomic) RNA synthesis

Our findings that reduced translational readthrough and expression of the alphavirus nsp4 lowers SINV fitness for non-opal stop codons appear to be at odds with our observation that sense-codon SINV variants exhibit lower fitness. Sense-codon variants should produce higher levels of nsP4 than opal since they do not depend on translational readthrough (Figure 1D, F, Suppl. Figure 2). However, higher nsP4 expression in sense-codon variants is offset by its rapid turnover via the ubiquitin-dependent N-end rule pathway ^29–31^. Thus, the mechanistic basis of fitness loss in sense-codon SINV variants must be fundamentally different than that for the ochre and amber SINV variants. We investigated the molecular basis of fitness loss in SINV sense-codon variants in Vero-37º.

Both P123 and P1234 polyproteins undergo proteolytic processing by the nsP2 protease (nsP2^Pro^) through a series of highly orchestrated steps (Figure 3A), as the nsP2^Pro^ sequentially transitions its cleavage site preference from P3/4 to P1/2 to P2/3 ^10^. Since sense-codon variants produce 85% more P3/4 substrate than WT SINV (Figure 2B), we hypothesized that excess P1234 production might delay this orderly transition, with deleterious consequences on viral fitness. To test this hypothesis, we conducted western blot analyses of Vero-37º cell lysates 18 hours post-infection to identify aberrations or delays in P1234 processing in sense-codon SINV variants (Figure 3B). We used a FLAG epitope-tagged nsP3 SINV strain, which allowed us to visualize nsP3 using an anti-FLAG antibody, as well as two polyclonal antibodies previously raised against SINV nsP2 and nsP4 proteins (Suppl. Figure 4A); no suitable anti-nsP1 antibody is available. Western blot analyses with all three reagents showed no discernible differences in the steady-state levels of fully processed nsP2 or nsP4 for any of the SINV strains passaged in Vero-37º (Figure 3B, Suppl. Figure 4A-B). In contrast, levels of nsP3 (which is heavily phosphorylated) were quantifiably lower in sense-codon SINV variants in Vero-37º cells (Suppl. Figure 4D). This indicates that the lower fitness of sense-codon variants is not caused by changes in protease (nsP2) or polymerase (nsP4) levels but might be correlated with lower levels of processed nsP3, consistent with previous reports ^12,30^.

**Figure 3.**
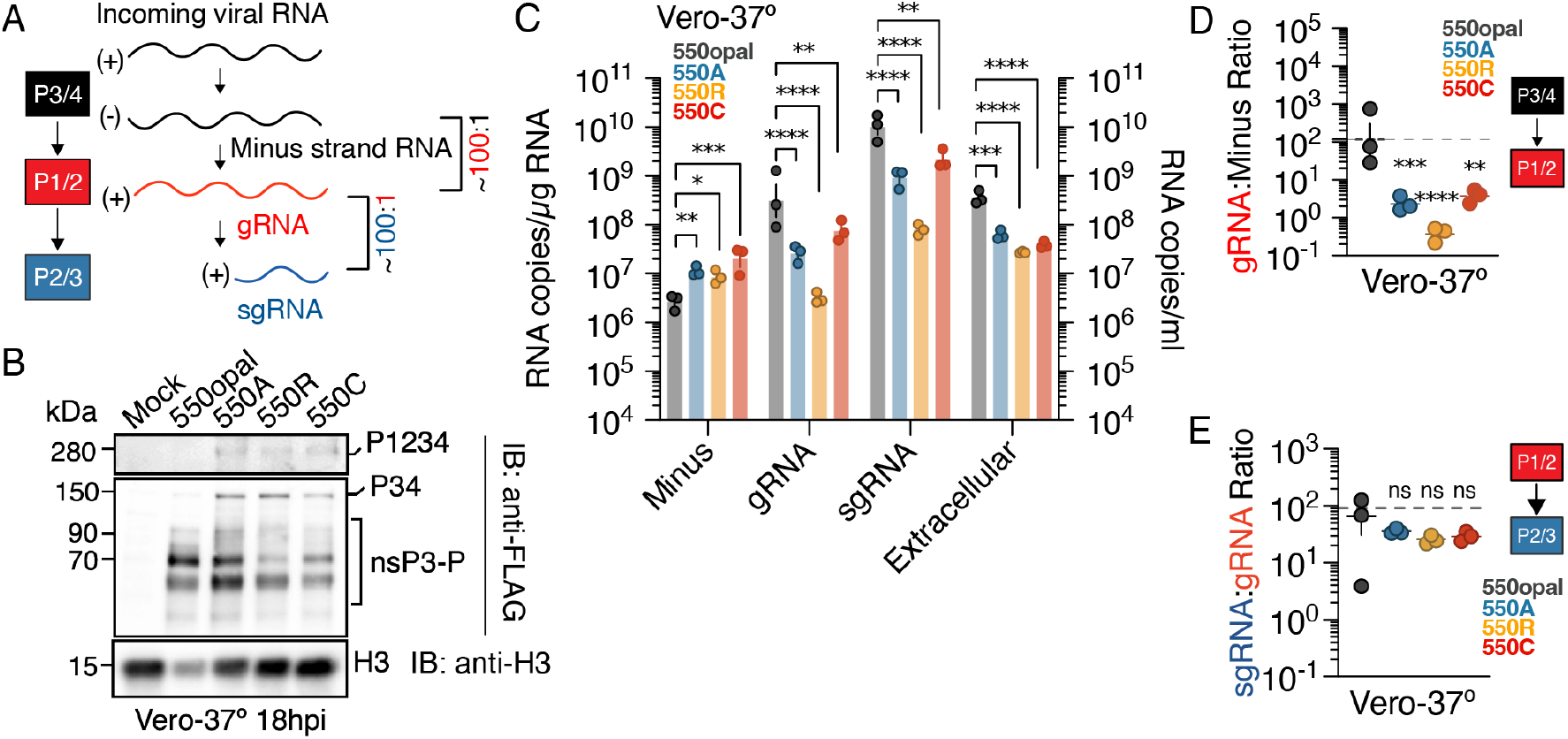
Aberrant polyprotein processing reduces the fitness of sense-codon SINV variants. (A) The current model of alphavirus nonstructural polyprotein processing summarizes the sequential processing steps catalyzed by the alphavirus nsP2 protease, which are required to form distinct processing intermediates that synthesize specific viral RNA species. Under optimal replication conditions, gRNA: minus-strand and sgRNA:gRNA ratios are expected to be 100:1. (B) Vero-37º cells were infected at MOI of 5 with WT (550opal) or sense-codon (550A, 550R, or 550C) SINV variants expressing 3XFLAG-tagged nsP3. 18h post-infection, total protein was extracted for western Blotting. Blots were probed with polyclonal sera against anti-FLAG and anti-histone H3 monoclonal antibodies. Data are representative of three independent experiments. (C) Effect of sense codon replacements on the synthesis of different viral RNA species in Vero-37º. SINV variants were used to infect Vero-37º at an MOI of 0.1 and harvested at 4- or 18-hours post-infection to quantify levels of minus-strand RNA, genomic RNA (gRNA), subgenomic RNA (sgRNA), or extracellular genomic RNA. (D) The ratio of SINV genomic (gRNA) to minus-RNA levels in Vero-37º (E) The ratio of subgenomic (sgRNA) to genomic (gRNA) RNA species in Vero-37º cells. Two-way ANOVA with Tukey’s multiple comparisons test. ^****^ = P < 0.0001, ^***^ = P < 0.001, ^**^ = P < 0.01, ^*^ = P < 0.05, ns = not significant.

Our western blot analyses also revealed significant evidence of delayed or aberrant polyprotein processing in Vero-37º cell lysates from sense-codon SINV infections relative to WT SINV. Using anti-FLAG and anti-nsP2 antibodies, we detected higher levels of the unprocessed P1234 polyprotein (280 kDa) in lysates from sense-codon SINV variants (Figure 3B, Suppl. Figure 4A). Additionally, we observed aberrant P12 and P34 products in the nsP2, nsP3, and nsP4 blots of sense-codon SINV variants. Quantifiably higher accumulation of P34 could be readily observed independently in the nsP3-FLAG and nsP4 blots (Figure 3B, Suppl. Figure 4E). Similarly, the presence of P12 in Vero-37º lysates suggests that P1/2 cleavage is also delayed or disrupted in sense-codon SINV variants (Suppl. Figure 4A). P12 and P34 intermediates can only be generated if the P2/3 cleavage aberrantly precedes the P3/4 and P1/2 cleavage steps (Figure 3B, Suppl. Figure 4A) ^32^. The high levels of P34 and P1234 in sense-codon SINV variants in Vero-37º contrast with the mostly undetectable levels of these aberrant products in WT SINV. They indicate delayed P3/4 cleavage despite the presence of proteases capable of cleaving this site. Collectively, our western blot analyses demonstrate nsP polyprotein processing defects among sense-codon SINV variants in Vero-37º.

Sequential and timely cleavage of the nsP polyprotein (P1234) is critical for synthesizing different viral RNA species in proportional amounts required for viral replication (Figure 3A) ^10,33^. Initial P1234 cleavage at the P3/4 site, located six amino acids downstream of the opal codon, gives rise to P123 and nsP4, constituting the replicase synthesizing the minus-strand RNA ^34^. The second cleavage reaction at the P1/2 site releases nsP1, P23, and nsP4, which causes an irreversible switch in RNA replication from minus-strand synthesis to plus-strand (genomic RNA, or gRNA) synthesis ^10,33,35,36^. The third and final cleavage reaction occurs at the P2/3 site, giving rise to fully processed nsP1, nsP2, nsP3, and nsP4 proteins, which together are responsible for producing predominantly sub-genomic RNA (or sgRNA) from a downstream promoter element on the minus strand RNA ^37^.

Since cleavage of the nsP polyprotein is so closely tied to changes in RNA synthesis from minus-strand to gRNA and from gRNA to sgRNA (Figure 3A), we next examined viral RNA levels using RT-qPCR in Vero-37º cells infected with either WT SINV or each of three sense-codon SINV variants (550A, 550R, or 550C) to infer how differences in polyprotein processing affected synthesis of different intracellular viral RNA species (Figure 3A). Four hours post-infection, we found that minus-strand levels were two to threefold higher than WT SINV for all three sense-codon SINV variants in Vero-37º. 18 hours post-infection, we found that gRNA levels were significantly lower for sense-codon SINV variants than for WT SINV (Figure 3C). As a result, while WT SINV (550opal) maintains the typical ratio of ∼100 gRNA copies per minus strand ^38,39^, sense-codon SINV variants have significantly lower gRNA, only making ∼1 gRNA copy per minus strand (Figure 3D). We also measured sgRNA levels in WT and sense-codon SINV variants to evaluate whether the processing defects persist or worsen in downstream steps. 18 hours post-infection, we found that sgRNA levels were also significantly lower in sense-codon SINV variants than in WT SINV (Figure 3C). However, all four SINV strains had comparable sgRNA: gRNA ratios of approximately 50 to 100, which are typical of WT SINV ^38,39^ (Figure 3E).

Our findings suggest that the transition from minus-strand to gRNA replication, which occurs upon P1/2 cleavage, is significantly delayed by the overproduction of the P1234 polyprotein in sense-codon SINV variants, resulting in higher minus-strand and much lower gRNA production. This reduction in gRNA production directly translates to fewer extracellular genome copies of SINV, *i*.*e*., lower virus production 24 hours post-infection in sense-codon SINV variants than WT SINV (Figure 3C). However, we infer that P2/3 cleavage is not delayed in the sense-codon SINV variants since sgRNA: gRNA ratios are normal (Figure 3G-H). Thus, our western blot analyses and RNA species profiling implicate aberrant polyprotein processing and a delayed switch from minus-strand to genomic RNA replication as the primary causes of the reduced fitness of sense-codon SINV variants in Vero-37º.

### Lower temperatures reduce the fitness advantage of opal-codon over sense-codon SINV variants

Previous studies have found that SINV replication kinetics is slower in mosquito cells than in vertebrate cells ^40,41^. To test whether the competitive fitness of sense-codon SINV variants is recovered at 28ºC (Figure 1F, Suppl. Figure 2) due to slower replication of WT SINV (550opal), we first measured replicative fitness of individual SINV variants across Vero-37º, Vero-28º and C6/36 cells. Indeed, by 24 hours, replication of WT SINV (550opal) in Vero-28º and C6/36 cells was approximately five-fold lower compared to Vero-37º (Figure 4A). In contrast, the replication of sense-codon variants was comparable across all three conditions. These results show that sense-codon variants can better compete against WT SINV at 28ºC because WT SINV replicates slower at this temperature. Indeed, we observed this trend of temperature-dependent tolerance of sense-codon variants (550A, 550R, 550C) across multiple vertebrate cell lines (see Suppl. Figure 5).

**Figure 4.**
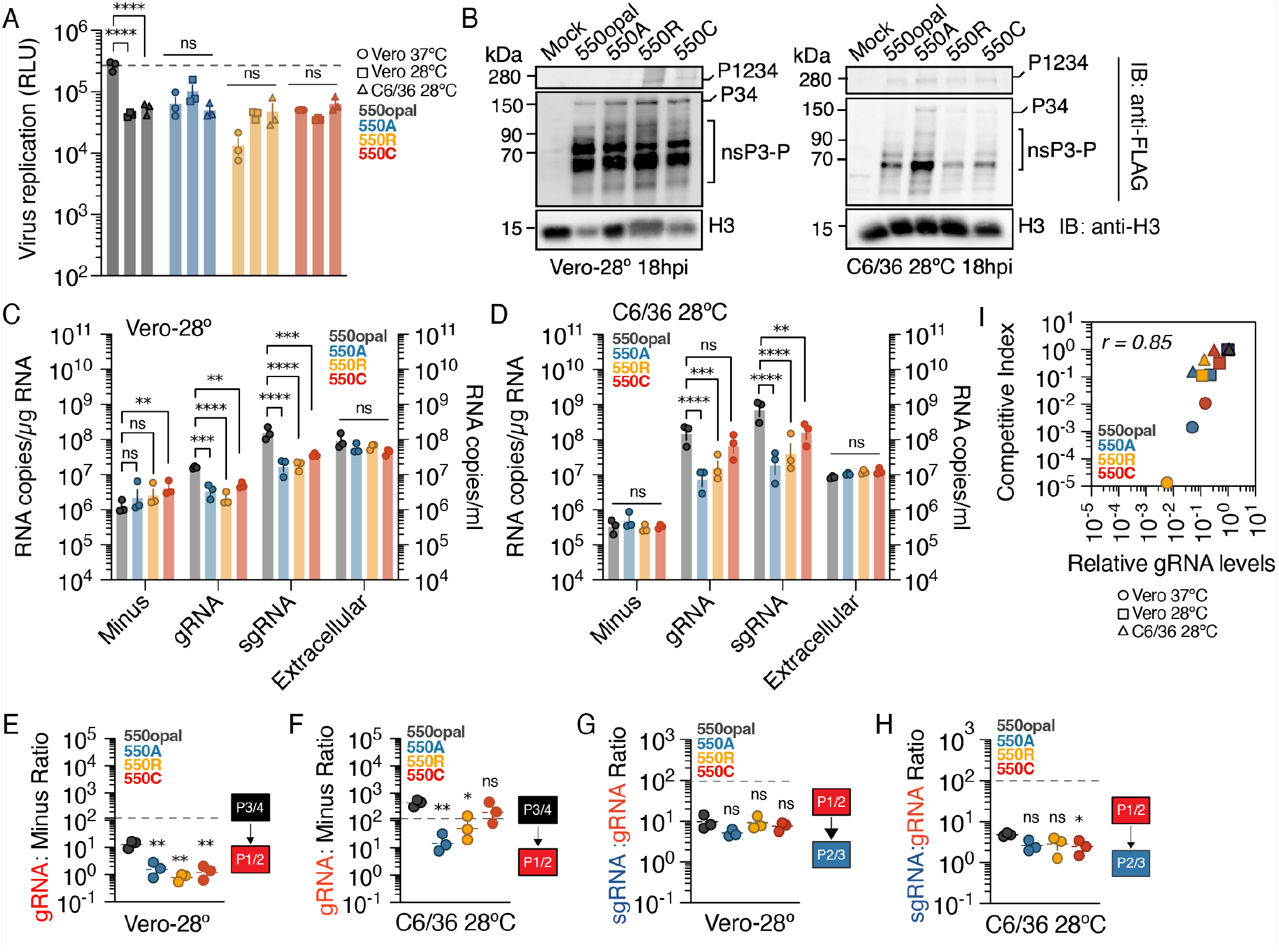
Fitness defects of SINV variants relative to WT SINV are reduced in mosquito cells. (A) A) Replication of WT (550opal) and sense-codon (550A, 550R, or 550C) SINV variants expressing translationally fused luciferase reporter in Vero-37º, Vero-28º and C6/36 cells. Cells were transfected with 100ng of capped, *in vitro* transcribed viral RNA and viral replication was measured 24-hours later using luciferase activity as a proxy for viral non-structural protein expression. (B) Vero-28º and C6/36 cells were infected at MOI of 5 with WT (550opal) or sense-codon (550A, 550R, or 550C) SINV variants expressing 3XFLAG-tagged nsP3. 18h post-infection, total protein was extracted for western blotting. Blots were probed with polyclonal sera against anti-FLAG and anti-histone H3 monoclonal antibodies. Data are representative of three independent experiments. (C-D) Effect of sense codon replacements on the synthesis of different SINV RNA species. Vero-28º (C) and C6/36 (D) cells were infected at an MOI of 0.1 and harvested at 4- and 18-hours post-infection to quantify levels of minus-strand, genomic (gRNA), subgenomic RNA (sgRNA), and extracellular genomic RNA. (E-F) Ratio of genomic (gRNA) to minus viral RNA species in Vero-28º (E) and C6/36 (F) cells infected with different sense codon variants. (G-H) Ratio of subgenomic (sgRNA) to genomic (gRNA) RNA species in Vero-28º (G) and C6/36 (H) cells infected with different sense codon variants. Two-way ANOVA with Tukey’s multiple comparisons test. ^****^ = P < 0.0001, ^***^ = P < 0.001, ^**^ = P < 0.01, ^*^ = P < 0.05, ns = not significant. (I) Pearson’s correlation between variant competitive index and relative gRNA levels. r^2^ = 0.73, 95% CI 0.5469 to 0.9580, P = 0.0004.

Western blot analyses also revealed only subtle differences between SINV wild-type (WT) and sense-codon SINV variants in Vero-28º (Figure 4B, Suppl. Figure 4B). We observed aberrantly processed P34 in all samples, with only slightly higher levels in sense-codon SINV variants compared to WT SINV (Figure 3B, Suppl. Figure 4B). This observation can be partly attributed to higher levels of translational readthrough at lower temperatures (see Figure 2B), presumably resulting in increased P1234 production even in WT SINV, accompanied by processing delays like in sense-codon variants. Furthermore, P12 was undetectable for WT SINV (550opal) and all sense-codon variants in Vero-28º, indicating at least a partial rescue of P1/2 processing defects observed in Vero-37º. Similarly, western blot analysis of infected C6/36 cell lysates showed either no (550C) or only subtle (550A, 550R) differences in nsP processing between WT SINV (550opal) and sense-codon SINV variants (Figure 4B, Suppl. Figure 4C).

Consistent with these western blotting results, we found that minus-strand RNA levels for 550A or 550R did not significantly differ from WT SINV in Vero-28º (Figure 4C); however, minus-strand levels were still higher in 550C compared to WT SINV. Moreover, gRNA and sgRNA levels were generally lower for all SINV variants, including WT SINV, in Vero-28º. Thus, all SINV variants exhibit low gRNA: minus-strand RNA and sgRNA: gRNA ratios in Vero-28º (Figure 4E,G). For example, WT SINV produces 10-fold fewer gRNA molecules per minus-strand RNA in Vero-28º than Vero-37º (Figure 3D, Figure 4C). Consequently, gRNA: minus-strand RNA ratios of sense-codon SINV variants are much less impaired (only 10-fold lower) than WT SINV in Vero-28º, whereas they are 100-fold lower in Vero-37º. Therefore, the replication of WT SINV is lowered to a degree such that it produces gRNA and extracellular virions at levels comparable to sense-codon variants in Vero-28º (Figure 4C).

In C6/36 cells, profiling of viral RNA levels revealed no differences between sense codon variants and WT SINV in minus-strand RNA levels of or sgRNA: gRNA ratios (Figure 4D). gRNA levels and gRNA: minus-strand RNA ratios were equivalent between the 550C variant and WT SINV, but both 550A and 550R showed significantly lower gRNA levels and gRNA: minus-strand RNA ratios than WT SINV (Figure 4F,H). However, unlike in Vero-37º or Vero-28º, even the impaired 550A and 550R SINV variants produce higher gRNA amounts than minus-strand RNA in C6/36 cells. Since gRNAs serve multiple roles during infection – from minus-strand RNA synthesis to nsP expression to virion production— gRNA levels are rate-limiting for viral fitness (Figure 1F, Suppl. Figure 2, Figure 4B-C). This enhancement of intracellular gRNA in C6/36 cells leads to higher extracellular genomic RNA in sense-codon variants, almost to the levels seen for WT SINV, which leads to comparable virus production across SINV variants in C6/36 cells (Figure 4B, I). These results can also explain why cysteine (550C) is the most tolerated amino acid substitution in C6/36 cells since it can produce wildtype levels of gRNA (Figure 1C, E-F, Suppl. Figure 2, Figure 4).

Our findings explain the higher tolerance of many sense-codon variants in Vero-28º and C6/36 cells. We find that higher translational readthrough and slower replication kinetics of WT SINV (550opal) in both Vero and C6/36 cells at 28ºC reduce the fitness gap between WT and sense-codon variants (Figure 1E-F, Figure 4A), resulting in near-equivalent levels of extracellular viral RNA (Figure 4C-D).

### A naturally co-occurring mutation can rescue 550C SINV variant fitness

In contrast to most SINV isolates found in nature, the neurovirulent AR86 strain naturally encodes a cysteine codon (UGC) instead of the opal stop codon ^42^. We hypothesized that one or more co-occurring mutations in SINV-AR86 might rescue the fitness of the 550C SINV variant in Vero-37º by restoring the proper processing of the nsP polyprotein P1234. Among several non-synonymous mutations in SINV-AR86, we were especially intrigued by the I538T mutation near the P1/2 cleavage site motif in nsP1 (Figure 5A). The I538T mutation has previously been shown to antagonize the JAK/STAT pathway, preventing the induction of Type I/II interferon responses, explaining why it may have been selected in vertebrate hosts ^43,44^. However, the I538T mutation also leads to delayed cleavage at the P1/2 site ^45^, which appeared to contradict our model because slower P1/2 cleavage would be predicted to exacerbate rather than ameliorate P1234 processing defects. However, slower P1/2 cleavage could also influence the efficiency of the downstream P2/3 cleavage. Indeed, previous studies have shown that uncleaved P123 cannot cleave a P23 substrate *in vitro* ^36^. Therefore, we hypothesized that introducing the I538T mutation into SINV sense-codon variants might restore proper nsP processing by decelerating both the P1/2 and P2/3 cleavage steps.

**Figure 5.**
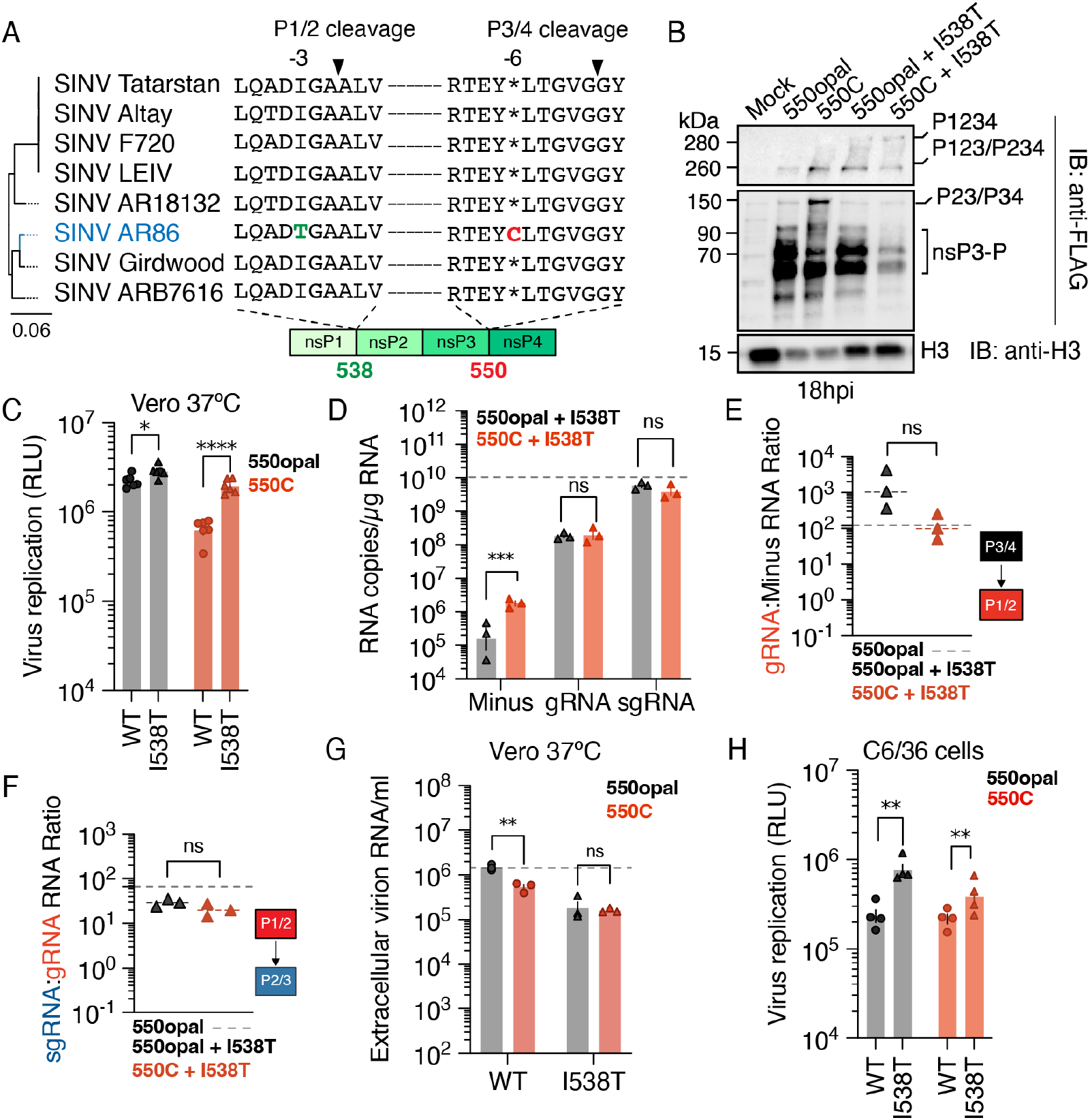
An nsP1 mutation can rescue the fitness defect of the 550C SINV variant in Vero-37º. (A) Multiple sequence alignment of the P1/2 and P3/4 cleavage sites in eight representative SINV strains (SINV AR86 is highlighted in blue). (B) WT (550opal) or 550C SINV variants expressing 3XFLAG-tagged nsP3, with or without the nsP1 I538T mutation, were used to infect Vero-37º cells at a MOI of 5. 18h post-infection, total protein was extracted for western blotting. Blots were probed with polyclonal sera against anti-FLAG and anti-histone H3 monoclonal antibodies. Data are representative of three independent experiments. (C) Replication of WT (550opal) or 550C SINV variants, with or without nsP1 I538T mutation, in Vero-37º. Cells were transfected with *in vitro* transcribed RNA of SINV variants viruses expressing an in-frame Nluc. Virus replication was assessed using luciferase assay 48h post-transfection. (D) Effect of I538T mutation on the synthesis of different viral RNA species. WT (550opal) or 550C SINV variants were used to infect Vero-37º at a MOI of 0.1 and harvested at 4- or 18-hours post-infection to quantify levels of minus-strand RNA, genomic RNA (gRNA), and subgenomic RNA (sgRNA). The horizontal dotted line denotes sgRNA copies produced by WT SINV (550opal). (E) Ratio of genomic (gRNA) to minus-strand viral RNA species in Vero-37º infected with WT (550opal) or 550C SINV variant carrying I538T mutation. (F) Ratio of subgenomic (sgRNA) to genomic (gRNA) viral RNA species in Vero-37º infected with WT (550opal) or 550C SINV variant carrying I538T mutation. The dotted lines represent ratios calculated for WT SINV (550opal). (G) Virus production resulting from infection of WT (550opal) or 550C SINV variants, with or without I538T mutation, in Vero-37º cells at an MOI of 0.1. Extracellular viral genome copies were quantified using RT-qPCR of supernatants collected 18h post-infection as a proxy for *de novo* virus production. (H) Replication of I538T variants in C6/36 cells, transfected with *in vitro* transcribed RNA of WT or I538T mutation-carrying SINV variants expressing an in-frame fused Nluc and either 550opal or 550C. Virus replication was assessed using luciferase assay 24h post-transfection. Two-way ANOVA with Tukey’s multiple comparisons test. ^****^ = P < 0.0001, ^***^ = P < 0.001, ^**^ = P < 0.01, ^*^ = P < 0.05, ns = not significant.

Western blot analyses of infected cell lysates collected 18 hours post-infection supported this ‘dual delay’ hypothesis, revealing the presence of P123/ P234 polyprotein in all variants, with higher levels observed in the 550C and Opal+I538T lysates (Figure 5B). This result confirms previous studies that reported a delay in P1/2 in Opal+I538T ^45^. We detected unprocessed P1234 in cell lysates infected with Opal+I538T and 550C+I538T, indicating that slowing down P1/2 cleavage also delays polyprotein processing at other sites (Figure 5B). The most compelling evidence of reduced P2/3 processing in the I538T mutants was the significantly reduced levels of the P34 polyprotein product in 550C+I538T compared to 550C (Figure 5B). Indeed, the levels of processed nsP3 are reduced considerably in the 550C+I538T variant compared to 550C, likely due to reduced P2/3 cleavage. In contrast, the levels of processed nsP3 remained unaltered in WT SINV, regardless of the presence of the I538T mutation (Figure 5B) ^45^. These results support our hypothesis that a double processing delay in the 550C+I538T variant restores viral fitness. Indeed, replication of 550C+I538T was significantly higher than 550C in Vero-37º and restored to the level of WT SINV (550opal) (Figure 5C).

Viral RNA profiling in Vero-37º cells revealed that 550C+I538T accumulated more minus-strand RNA than 550opal+I538T (Figure 5D). In contrast, both variants produced equivalent levels of plus-strand gRNA and sgRNA (Figure 5D). In addition, the gRNA to minus-strand RNA ratio in 550C+I538T infected cells was restored to WT SINV (550opal) level (Figure 5E, Figure 3E). Consistent with our hypothesis that I538T mutants delay the switch from gRNA to sgRNA production, we found that both 550opal+I538T and 550C+I538T mutants produced less sgRNA compared to WT SINV (Figure 5D). This is also reflected in lower sgRNA to gRNA ratios for 550opal+I538T and 550C+I538T compared to WT SINV (550opal) (Figure 5F, Figure 3G). Thus, 550C SINV variants compensate for a delayed transition in gRNA production by delaying the transition from gRNA to sgRNA production, enabling sufficient time for gRNA production. Reduced sgRNA levels in I538T mutants are not inconsequential; they lead to lower virus production, likely due to insufficient expression of structural proteins. As a result, although both 550opal+I538T and 550C+I538T SINV variants produced equivalent amounts of virus 18 hours post-infection, they still produce less virus than either WT or 550C SINV variants (Figure 5G). Finally, in C6/36 cells, 550opal+I538T and 550C+I538T replicated better than their non-I538T counterparts (Figure 5H), suggesting that delayed P123 processing increases SINV replication in mosquito cells. These findings suggest that sense-codon SINV variants have selective advantages that allow them to thrive in mosquito vectors but are disadvantaged in vertebrate cells unless they acquire compensating mutations like I538T. Thus, our study unravels the mechanistic basis for the host (temperature)-specific selective constraint that leads to the retention of the opal codon in most dual-host alphaviruses.

## Discussion

In this study, we investigated the biological cause underlying the widespread retention of the premature opal termination codon in nsP3 across alphaviruses. Using saturation mutagenesis and pooled selection, we comprehensively assessed all possible substitutions at a single codon in the prototype alphavirus SINV. These and subsequent experiments led us to conclude that host temperature is the predominant source of selective constraint at this site (Figure 1), with the opal codon being significantly preferred over all others in Vero-37º. However, this preference is significantly lower at 28º in primate and mosquito cells. In Vero-37º, the opal termination codon ensures the optimal degree of readthrough that produces sufficient levels of the nsP4 viral polymerase, which other stop codons cannot (Figure 2), without producing so much P1234 polyprotein, as sense-codon variants do, to disrupt its proteolytic processing cadence (Figure 3-4). Lower temperatures in vertebrate or mosquito cells increase translational readthrough (restoring fitness of alternate stop codons) and lead to similar processing delays in WT and sense-codon variants, reducing their fitness gap (Figure 3-4). Unexpectedly, we find that nsP processing defect in a sense-codon SINV variant in Vero-37º can be compensated by an additional delay in a subsequent step, which helps restore the proper orchestration of nsP polyprotein processing (Figure 5). However, consistent with previous reports, such delays in polyprotein processing may actually benefit alphavirus replication in mosquito cells (Figure 5H) ^46^.

Our findings provide insights into how alphaviruses encoding opal to sense codon mutations exist and may even thrive in nature. Cysteine variants (the only naturally occurring opal-to-sense mutation found in SINV) exhibit fitness levels comparable to or exceeding those of WT SINV in mosquito cells. Another alphavirus, EEEV, also acquired opal-to-cysteine mutations upon passaging in mosquito cells^14^. Therefore, we propose that cysteine mutations spontaneously arise and are favored in alphaviruses in the mosquito vector. Our results also demonstrate why the co-occurring I538T mutation from SINV-AR86, which blunts the Type I/II interferon responses, ^**43**,**44**^ is essential for the cysteine mutation’s viability in vertebrate cells at 37ºC. Therefore, the I538T and 550C mutations might co-occur to confer fitness advantages in both vertebrate and mosquito cells (Figure 5C, H). Future studies could evaluate whether other dual-host alphaviruses, particularly those within the Semliki Forest Virus (SFV) clade, use analogous strategies to tolerate the presence of sense codons at the opal codon site.

Temperature is a critical environmental cue sensed by many microbial pathogens, including viruses, that navigate through multiple hosts, enabling them to adapt their replicative strategies to their new host environment instantaneously ^47–49^. Arboviruses like West Nile (WNV) have been shown to encode molecular mechanisms that sense shifts in host temperature and adjust their replicative strategies ^48^. Temperature is especially critical for RNA viruses like alphaviruses as it can shape many fundamental aspects of their genome replication, transcription, and translation ^33,49^. Our findings suggest that the nsP3 opal codon adapts the replication of dual-host alphaviruses like SINV to host temperatures. Indeed, phylogenetic analysis of alphaviruses shows a correlation between the presence of the nsP3 opal codon and the alphaviruses’ ability to infect endothermic hosts or at high ambient temperatures (Suppl. Figure 6). In contrast to the strong conservation of the opal codon in dual-host alphaviruses that navigate through insect and vertebrate hosts, some alphaviruses that infect marine vertebrates at near-freezing temperatures encode a sense codon (glutamine: CAA, CAG) instead. It is likely that the extremely high readthrough efficiency of the opal codon at low temperatures renders it unnecessary at the near-freezing temperatures encountered by its marine vertebrate hosts ^50,51^. However, marine alphaviruses that infect endothermic mammals, *e*.*g*., seals (Southern Elephant Seal Virus, SESV) and porpoises (Harbor Porpoise Alphavirus, HPAV) or tropical fish species (Wenling Alphavirus, WEAV), still preserve the opal codon, presumably because it still serves a useful role at higher temperatures (Suppl. Figure 6). Since marine alphaviruses are considered an ancestrally branching clade, the nsP3 opal codon thus represents an early innovation in alphavirus evolution ^**52**^. However, our hypothesis of temperature-dependent retention of the opal codon does not explain the near-universal preservation of the opal codon even in insect-restricted alphaviruses, suggesting additional insect host-specific selective constraints might exist in this alphaviral lineage. Indeed, although our *in vitro* model reveals mechanistic insights into the mutational constraints affecting the nsp3 opal codon, it may not fully account for additional constraints that could be present *in vivo* within vertebrate or mosquito host species ^**53**,**54**^.

The use of premature termination codons is widespread among RNA viruses. In particular, translational readthrough mechanisms are present outside of alphaviruses within some members of the alpha-like supergroup, including several plant viruses like tobraviruses, tobamoviruses, and furoviruses ^8^. These viruses also employ PRT of “leaky” termination codons to synthesize their viral replicases ^**28**,**55**^. Our work suggests that the opal codon serves as a crude but effective thermometer for viral replication, indicating that deploying leaky termination codons might be a broad strategy to sense and respond to temperature stress, whether that is abiotic in the case of plant viruses and alphaviruses that infect ectothermic hosts, or biotic, in the case of host-switching alphaviruses infecting endothermic vertebrates.

## Acknowledgements

We thank Tiia Freeman, Nandan Gokhale, Jennifer Hyde, Grant King, Irene Newton, Maria Toro Moreno, Jeannette Tenthorey, and Maria Jose Olmo Uceda for providing valuable feedback on the manuscript. We also thank members of the Malik and Emerman labs for providing intellectually stimulating discussions. We thank Richard Hardy and Suchetana (Tuli) Mukhopadhyay (Indiana University Bloomington) and Diane Griffin and the Griffin lab (Johns Hopkins University) for sharing valuable resources, including the parental SINV infectious clones and anti-nsP2 and anti-nsP4 antibodies. We thank the Hutch Shared Resources, especially the Genomics and Flow Cytometry Core Facilities, for their support. This work was supported by a National Institutes of Health grant (U54 AI170792 (PI: Nevan Krogan) to ME, HSM), Helen Hay Whitney Fellowship (TB), and a Howard Hughes Medical Institute Investigator award (to HSM). Funding agencies had no role to play in the execution of the project or the decision to publish. This paper was typeset with the bioRxiv word template by @Chrelli: www.github.com/chrelli/bioRxiv-word-template.

## Author contributions

Conceptualization: TB, ME, HSM

Methodology: TB, EMA, HSM

Investigation: TB, EMA, ACN

Visualization: TB

Funding acquisition: TB, ME, HSM

Project administration: TB, HSM

Supervision: ME, HSM

Writing – original draft: TB, EMA, ME, HSM

Writing – review & editing: TB, EMA, ACN, ME, HSM

## Materials and Methods

### Insect and Mammalian Cell Culture

C6/36 *Aedes albopictus* cells were grown at 28ºC under 5% CO_2_ in humidified incubators and were cultured in high-glucose, L-glutamine Minimal Essential Medium (Gibco) supplemented with 10% fetal bovine serum (Cytiva) and 1% penicillin-streptomycin (Gibco). African Green Monkey (*Chlorocebus aethiops*) - derived Vero and BSC40 cells, Human Embryonic Kidney cells (HEK293T), and Human hepatocyte epithelial (Huh7) cells were grown at 37ºC under 5% CO_2_ in humidified incubators. They were cultured in high-glucose, L-glutamine Minimal Essential Medium (Gibco) supplemented with 10% fetal bovine serum (Cytiva) and 1% penicillin-streptomycin (Gibco).

### Saturation Mutagenesis Screen

The Opal variant library was generated via PCR mutagenesis. Briefly, a 3.8 kb template spanning P34 was PCR amplified from the wildtype TE3’2J: GFP infectious clone (Fwd: 5’-GTATCG-TACTTACCGGTTGCCAGTGGAGC-3’, Rev: 5’-CAC-TCGATCAAGTCGAGTAGTGGTTGATC-3’). Mutagenic primers carrying ambiguous bases (NNN) at the opal codon site were used to generate the insert for downstream Gibson assembly. The TE3’2J: GFP viral backbone fragment lacking wildtype nsP3 was generated by digesting the infectious clone with AgeI and BstEII enzymes. Backbone and insert components were gel purified, mixed in 1:3 molar ratios, assembled via a two-part Gibson assembly (NEB HiFi assembly mix), and transformed into Endura DUO Electrocompetent Cells (LGC Bioresearch Technologies). Plasmid DNA was recovered from approximately 70,000 colonies collected and pooled from three independent replicates to ensure ∼1,000-fold coverage of the library of 64 variants. The entire screen was performed twice independently. Two µg of XhoI-linearized variant library DNA or WT TE3’2J: GFP was used for *in vitro* transcription reaction using SP6 RNA polymerase to generate pooled, capped viral RNA. *In vitro*, transcribed viral RNA was transfected into approximately two million cells using Lipofectamine LTX according to the manufacturer’s protocol (Thermo-Fisher). Viral infection rates were assessed by monitoring GFP fluorescence, and supernatants were collected at 48hpi in Vero cells grown at 37ºC and 28ºC and 72hpi in C6/36 cells grown at 28ºC. The sequencing library was generated as follows: Virion RNA was isolated from collected supernatants using the QIAMP viral RNA isolation kit (Qiagen), reverse transcribed using Superscript III Reverse Transcriptase (Invitrogen), and a SINV-specific RT primer (5’-GTCGGATGA-TATTTCTCCAAAGGCGC-3’). Purified cDNA from post-selection virion RNA or pre-selection library plasmid DNA was used to amplify the entire AgeI/BstEII flanked 3.8 kb region spanning the nsP3-nsP4 using KOD Hot-start Master Mix and the following cycling conditions: 1. 95ºC 2 min, 2. 95ºC 20 sec, 3. 70ºC 1 sec, (cool at 0.5ºC/s) 4. 50ºC 30 sec, 5. 70ºC 40 sec (Repeat steps 2-5 24x) 4ºC hold. This PCR (PCR0) product was purified using AMPure beads and diluted to 0.5ng/µl before generating UMI (Unique Molecular Identifier)-tagged amplicons spanning the opal codon site. The UMI-tagging PCR1 using KOD Hot-start Master Mix was carried out with the following cycling conditions: 1. 95ºC 2 min, 2. 95ºC 20 sec, 3. 70ºC 1 sec, (cool at 0.5ºC/s) 4. 50ºC 20 sec, 5. 70ºC 20 sec (Repeat steps 2-5 nine times) 6. 95ºC 1 min, 4ºC hold. UMI-tagged PCR1 was AMPure bead-purified and diluted to bottleneck to 3E5 barcodes, which corresponds to ≥ 3 reads/barcode at the target sequencing read depth. The final PCR2 step of the sequencing library preparation stage involved amplification using indexing primers with the following cycling condition: 1. 95ºC 2 min, 2. 95ºC 20 sec, 3. 70ºC 1 sec, (cool at 0.5ºC/s) 4. 55ºC 20 sec, 5. 70ºC 20 sec (Repeat steps 2-5 23x) 4ºC hold. Finally, all indexed PCR2 products were AMPure bead-purified before and after gel purification and pooled before sample submission for next-generation sequencing. Sequencing was performed on an Illumina MiSeq Nano V3 at PE150, which generated approximately 21 million paired-end reads. Codon counts for each variant were determined from reads with sequence quality ≥ Q30 using the dmstools2 package ^56^. Selection scores for individual codon variants were calculated as the ratios of variant codon counts post-selection vs codon counts pre-selection input library divided by the ratio of wildtype codon counts post-selection vs pre-selection input library.

### Virus Constructs

All viral stop codon mutant plasmids were generated via sitedirected mutagenesis. Mutagenic primers were designed using Takara’s In-Fusion Cloning Primer Design tool. PCR reactions were performed using Phusion polymerase (NEB) with the following cycling conditions: 1. 98ºC for 30 sec 2. 98ºC 30 sec 3. 50ºC for 30C 4. 72ºC 7 min (Repeat steps 2-4 18x) 5. 72ºC for 10 min 6, Hold at 4ºC. Reactions were then DpnI-treated for 6 hours and purified using the Monarch PCR and DNA Cleanup Kit (NEB) according to the manufacturer’s protocol. The purified reaction was then transformed into DH5α cells, and DNA was isolated using the Monarch Plasmid Miniprep Kit (NEB) according to the manufacturer’s protocol. (see Suppl Table 1 for specific protocols). Translation and translational readthrough reporters were generated by digesting parental and mutant plasmids and cloning in luciferase (Nano/Firefly) using Gibson assembly (NEB). The Gibson reactions were then transformed into DH5α cells, and DNA was isolated using the Monarch Plasmid Miniprep Kit (NEB) according to the manufacturer’s protocol. (see Suppl. Table 1 for specific protocols and primers).

### Independent Viral Growth Assays

2µg infectious clones were linearized with XhoI (NEB) and subjected to in-vitro transcription (IVT) using SP6 RNA polymerase (NEB) according to the manufacturer’s suggested protocol. C6/36 and Vero cells were seeded into 24-well to 70-80% confluency and transfected with IVTs using Lipofectamine LTX (Thermo-Fisher) according to the manufacturer’s protocol. Cells were collected 48 hours post-transfection and analyzed by flow cytometry (BD Fortessa).

### Viral Competition Assays

Genome copies of WT and mutant virus stocks were calculated using quantitative RT-PCR. Briefly, cDNA synthesis was performed on viral supernatant using M-MuLV Reverse Transcriptase (NEB) with oligo(dT) (20mer+5’Phos) (IDT) according to the manufacturer’s protocol. RT-qPCR analysis was performed using SYBR Green master mix (Thermo-Fisher) with gene-specific primers according to the manufacturer’s protocol and the Applied Bioscience StepOnePlus-qRT-PCR machine (Life technologies). Once obtained, WT and mutant SINV virus stocks were normalized to equal genome copies. C6/36 and Vero cells were seeded into 24 wells and infected with TE3’2J-mCherry wildtype virus alongside TE3’2J: GFP wildtype or mutant viruses at a 1:1 ratio (MOI – 0.1). Cells were collected 48 hours post-infection and analyzed by flow cytometry (BD Fortessa) to determine the percentage of cells infected with wildtype (red) and variant (green) viruses.

### Total RNA Extractions and Real-Time Quantitative RT-PCR Analysis

To quantify minus-strand RNA copies, cells were seeded into a 24-well and infected with WT or variants at >90% confluency at an MOI = 5. At 4 hours post-infection, cells were harvested using TRIZOL reagent (Thermo-Fisher), and RNA was extracted using the Direct-zol RNA Miniprep kit (Zymo) according to the manufacturer’s protocol. Following RNA extraction, cDNA was synthesized using M-MuLV Reverse Transcriptase (NEB) with minus-strand specific primers (IDT). To quantify plus-strand RNA copies, cells were seeded into a 24-well and infected with WT or variant SINV at >90% confluency at an MOI = 0.1. Cells were then harvested 24 hours post-infection using TRIZOL reagent (Thermo-Fisher). RNA was extracted using the Direct-zol RNA Miniprep kit (Zymo) according to the manufacturer’s protocol. Following RNA extraction, cDNA was synthesized using M-MuLV Reverse Transcriptase (NEB) with gene-specific primers and OligoDT (20mer+5’Phos) (IDT) according to the manufacturer’s protocol. In all cases, RT-qPCR analysis was performed using SYBR Green master mix (Thermo-Fisher) with plus-strand specific primers according to the manufacturer’s protocol and using the Applied Bioscience StepOnePlus-qRT-PCR machine (Life technologies).

### Viral Translation Assays

SINV nsP3-Nluc reporter viruses (Suppl. Table 1) were generated by digesting 2µg plasmid DNA with XhoI and performing in-vitro transcription (IVT) using SP6 RNA polymerase (NEB) according to the manufacturer’s protocol. C6/36 and Vero cells were seeded into black-welled, clear-bottom 96-well plates at 70-80% confluency and transfected with 100ng capped viral RNA using Lipofectamine LTX (Thermo-Fisher) according to the manufacturer’s protocol. At 24-hours post-transfection, translation was quantified using the NanoGlo luciferase assay system (Promega) according to the manufacturer’s protocol. Luminescence was recorded using a Cytation3 Imaging Reader (Bio-Tek).

### Translational Readthrough Assay

C6/36 and Vero cells were seeded into 96 wells at 70-80% confluency and transfected with the dual-reporter translational readthrough constructs (Suppl. Table 1) using Lipofectamine LTX (Thermo-Fisher) according to the manufacturer’s protocol. At 48 hours post-transfection, Fluc and Nluc signals were quantified using the Dual-Glo luciferase assay system (Promega) according to the manufacturer’s protocol. Luminescence was recorded using a Cytation3 Imaging Reader (BioTek). Fluc to Nluc ratios obtained for each stop codon variant were compared to a control construct carrying a sense mutation in place of the opal, representing a 100% translational readthrough.

### Western Blot Analyses

C6/36 and Vero cells were seeded into six-well plates and infected at >90% confluency (MOI = 5). Cells were washed with cold 1X PBS and harvested using RIPA buffer (Pierce) supplemented with 1X protease inhibitor (cOmplete). Purified protein was denatured using 2X Laemmli buffer (BioRad) with 5% β-Mercaptoethanol (Sigma-Aldrich). Western blots were run on Mini-PROTEAN precast gels and then transferred to a Trans-Blot 0.2 µm nitrocellulose membrane using the Trans-Blot Turbo Transfer System (Bio-Rad). Blots were incubated in 5% BSA and either primary α-FLAG antibody (Proteintech) at 1:3,000, α-nsP2 antisera at 1:3000, α-nsP4 antisera at 1:2000 or α-H3 antibody (Abcam) at 1:3,000 dilution overnight at 4ºC and washed with 1X TBS with 0.1% Tween-20. The blots were then probed with secondary α-rabbit HRP conjugate (R&D systems). Blots were visualized using a BioRad ChemiDoc Imaging System. Western blots probing for nonstructural protein expression and polyprotein processing were performed with independent biological replicates for each virus under different host conditions.

### Phylogenetics and Sequence Analysis

A multiple sequence alignment of nsP4 coding sequences from 49 extant alphavirus species was constructed using Clustal Omega. Alignments were manually curated using Geneious and maximum likelihood phylogenetic trees were generated using the HKY85 substitution model in PHYML, using 100 bootstrap replicates for statistical support. Phylogenetic trees were visualized using FigTree. Logoplots were generated using Skylign ^57^. Transcriptome-weighted codon usage counts for African green monkey (*Chlorocebus aethiops*) and Asian tiger mosquito (*Aedes albopictus*) derived from available GenBank and RefSeq data were obtained from CoCoPUT (https://dnahive.fda.gov/) ^23,24^.

### Statistics

Statistical analyses were conducted using GraphPad Prism (v10.2, GraphPad Software Inc., San Diego, CA). Data were first checked for a normal (Gaussian) distribution using the Kolmogorov-Smirnov test. For data meeting normality (α = 0.05) and similarity of variance criteria, means were compared using Twoway ANOVA (for multiple groups with two variables) with post hoc tests for multiple comparisons.

---

**Suppl. Figure 1.**
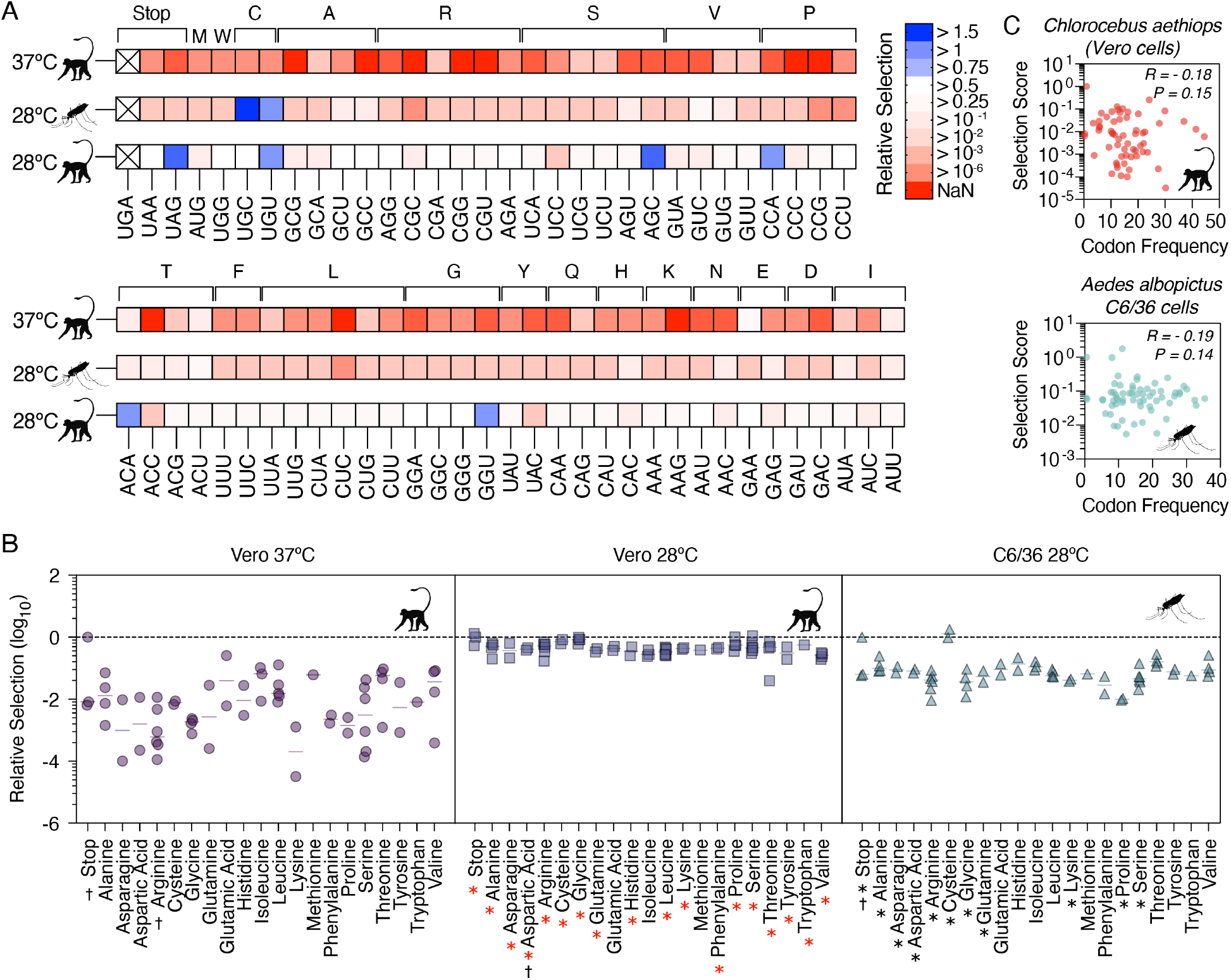
Selection scores of codon variants across cell types and temperatures. (A) Heat-map of selection scores of all codon variants, grouped based on encoded amino acids (on top). (B) Mean selection scores of SINV variants encoding different amino acids. Dotted line represents the selection score of WT (opal) SINV. Each data point is a synonymous codon encoding the corresponding amino acid. † denotes statistically significant different selection scores between synonymous codons in the same condition. Red asterisks indicate statistically significant differences between Vero-37º and Vero-28º cells, whereas black asterisks indicate statistically significant differences between Vero-37º and C6/36 cells. Two-way ANOVA with Tukey’s test for multiple comparisons. Pairwise unpaired t-tests with Welch’s correction. (C) Pearson’s correlation between selection scores and codon usage frequencies in Vero and C6/36 cells.

**Suppl. Figure 2.**
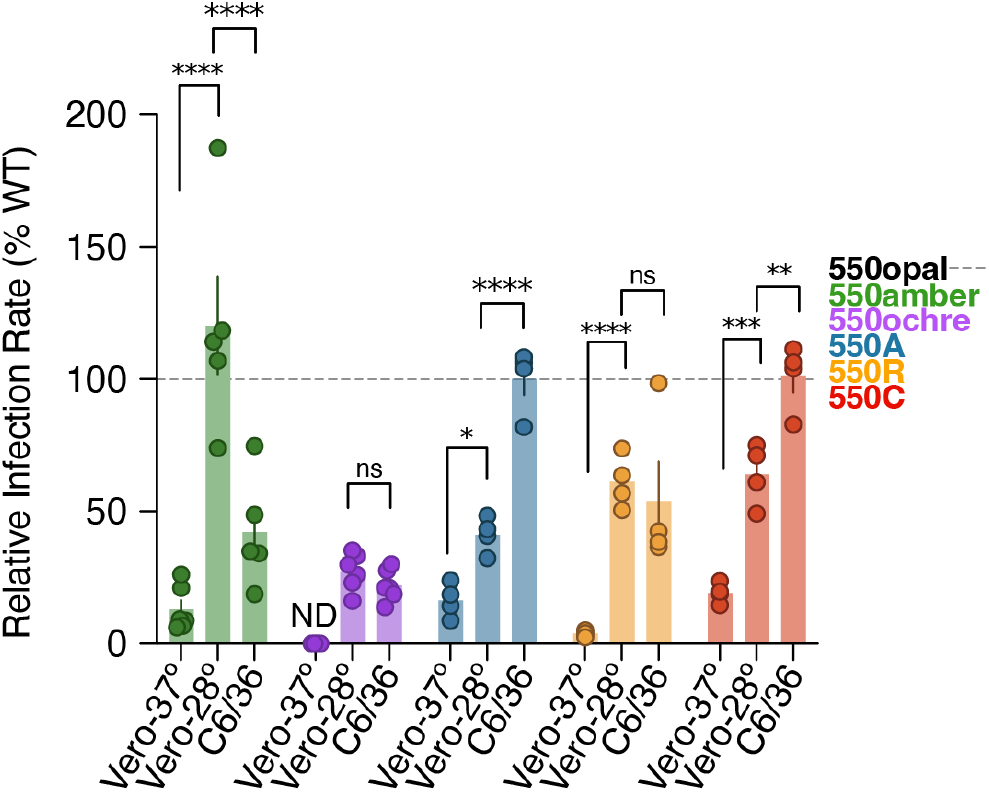
Fitness of alternate stop and sense-codon SINV variants. Alternate-stop codon variants (550amber and 550ochre) and sense-codon variants (550A, 550R, 550C) were engineered into a SINV-GFP reporter virus. Independent growth of variants was tested by infecting Vero-37º, Vero-28º, or C6/36 cells at MOI of 0.1 and quantifying GFP-expressing cells via flow cytometry 48 hours post-infection. Relative infection rates are normalized to WT SINV (550opal). Two-way ANOVA with Tukey’s multiple comparisons test. ^****^ = P < 0.0001, ^***^ = P < 0.001, ^**^ = P < 0.01, ^*^ = P < 0.05, ns = not significant, ND – not detectable.

**Suppl. Figure 3.**
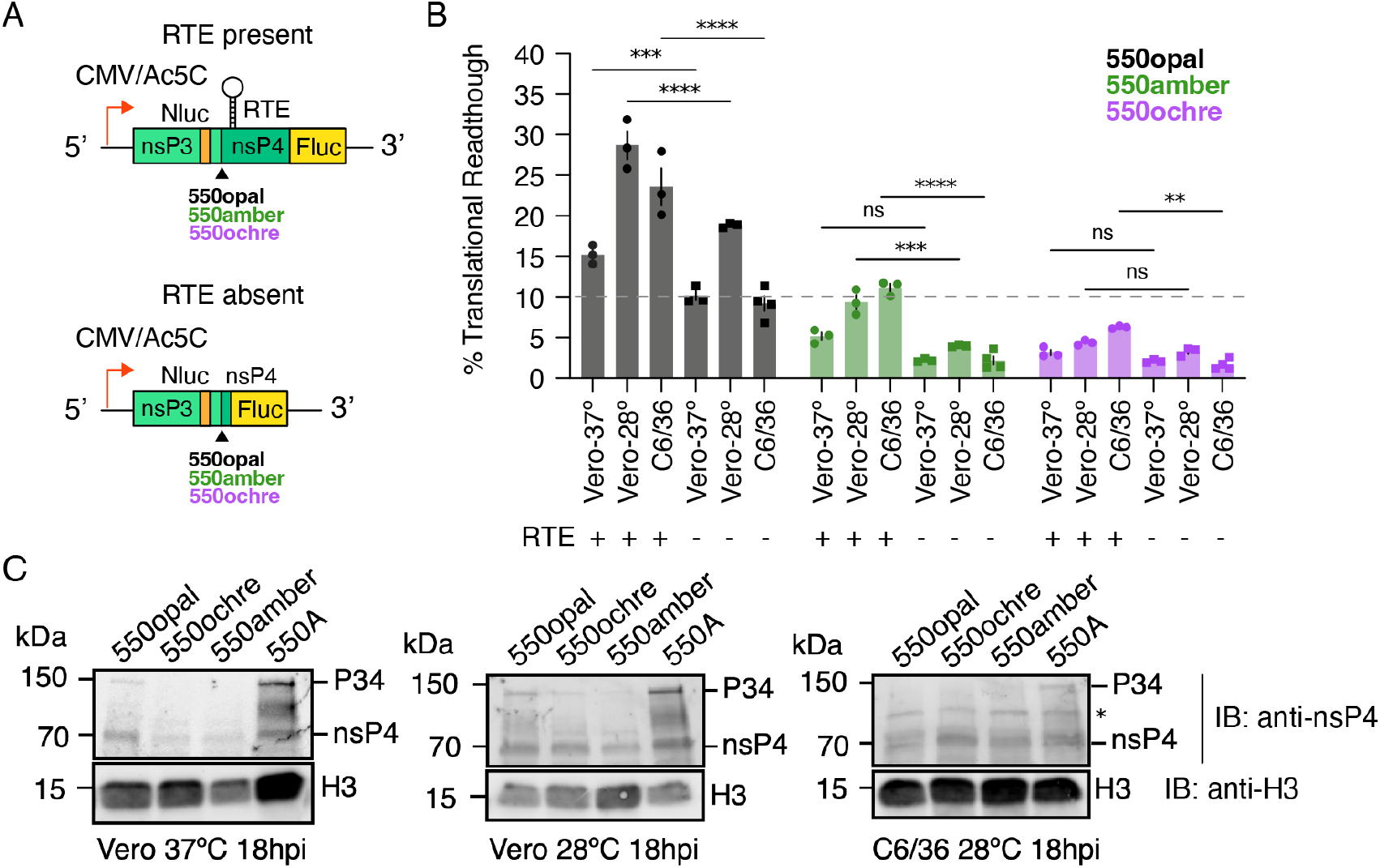
The termination codon <u>r</u>ead<u>t</u>hrough <u>e</u>lement (RTE) is required for the translational readthrough in vertebrate and mosquito cells. (A) Translational readthrough constructs were designed to include (top) or exclude (bottom) the downstream RNA structure, known as the termination codon <u>r</u>ead<u>t</u>hrough <u>e</u>lement (RTE). (B) Using reporters, translational readthrough was quantified relative to a reporter encoding a sense codon (GCA), which does not require translational readthrough for nsP4 expression. The horizontal dotted line denotes a 10% translational readthrough. Two-way ANOVA with Sidak’s multiple comparisons test. (C) Vero-37º, Vero-28º, or C6/36 cells were infected at MOI of 5 with WT (550opal), 550amber, 550ochre, or 550A sense-codon SINV variants. 18h post-infection, total protein was extracted for western blotting. Blots were probed with polyclonal sera against anti-nsP4 and anti-histone H3 monoclonal antibodies (as loading controls). Lane denoted with asterisk (*) represents a non-specific band observed in C6/36 cell lysates probed with anti-nsP4 polyclonal antibody.

**Suppl. Figure 4.**
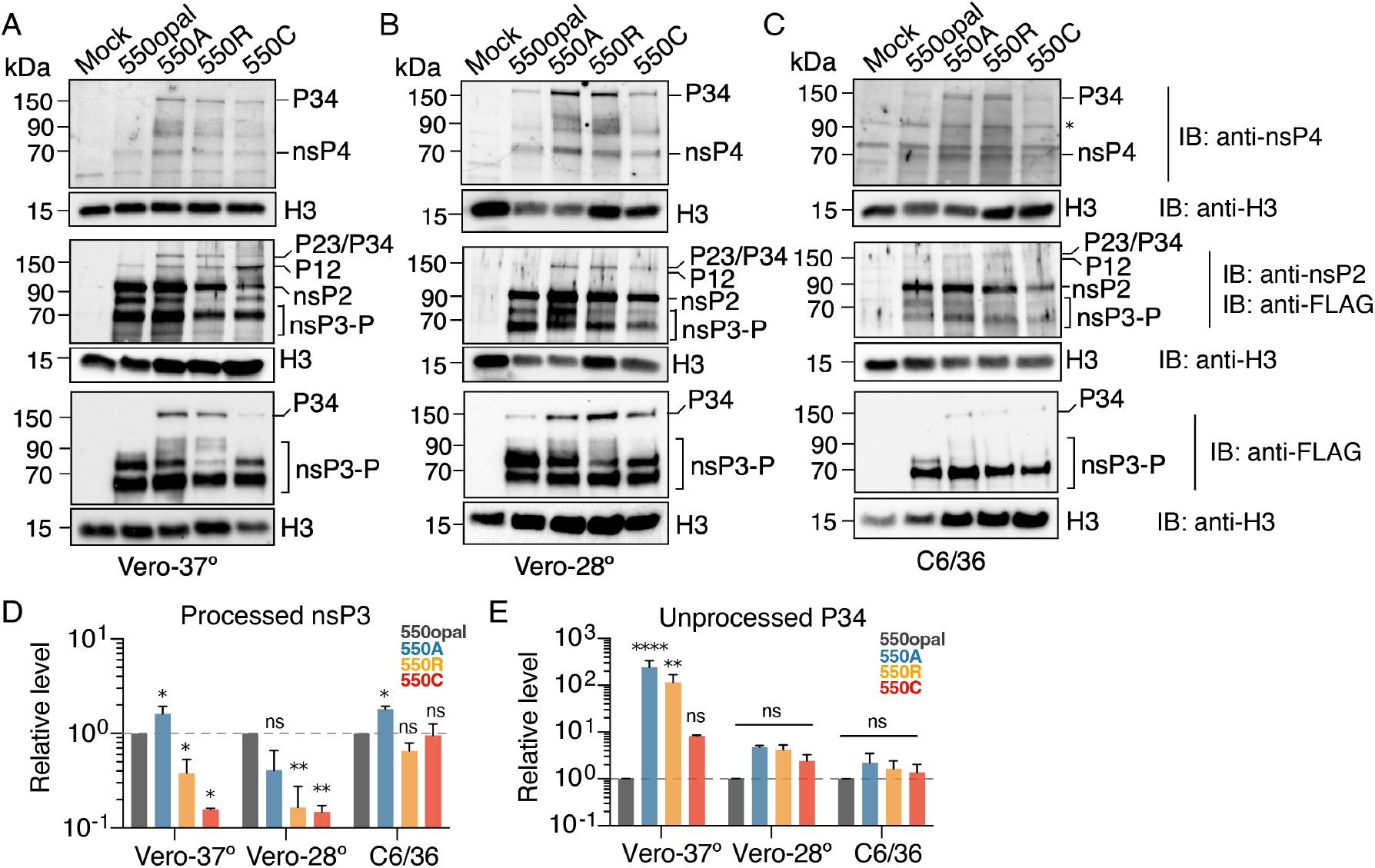
The presence of sense codons at the nsP3 opal codon site impairs alphavirus nonstructural polyprotein processing. (A) WT (550opal) or sense-codon (550A, 550R, or 550C) SINV expressing 3XFLAG-tagged nsP3 were used to infect Vero-37º cells. 18h post-infection, proteins were extracted and analyzed via western blotting using polyclonal sera against nsP2 and nsP4, anti-FLAG, and anti-histone H3 monoclonal antibodies. Data are representative of three independent experiments. Lane denoted with asterisk (*) represents a non-specific band observed in C6/36 cell lysates probed with anti-nsP4 polyclonal antibody. We also analyzed SINV-infected Vero-28º (B) or C6/36 (C) cells using the same protocols. Using ImageJ software, we quantified the bands observed in our western blotting analyses through densitometric analyses for nsP3-FLAG (D) or unprocessed P34-FLAG (E). In both cases, band intensities were normalized using H3. Data are representative of three independent experiments. Two-way ANOVA with Sidak’s multiple comparisons test.

**Suppl. Figure 5.**
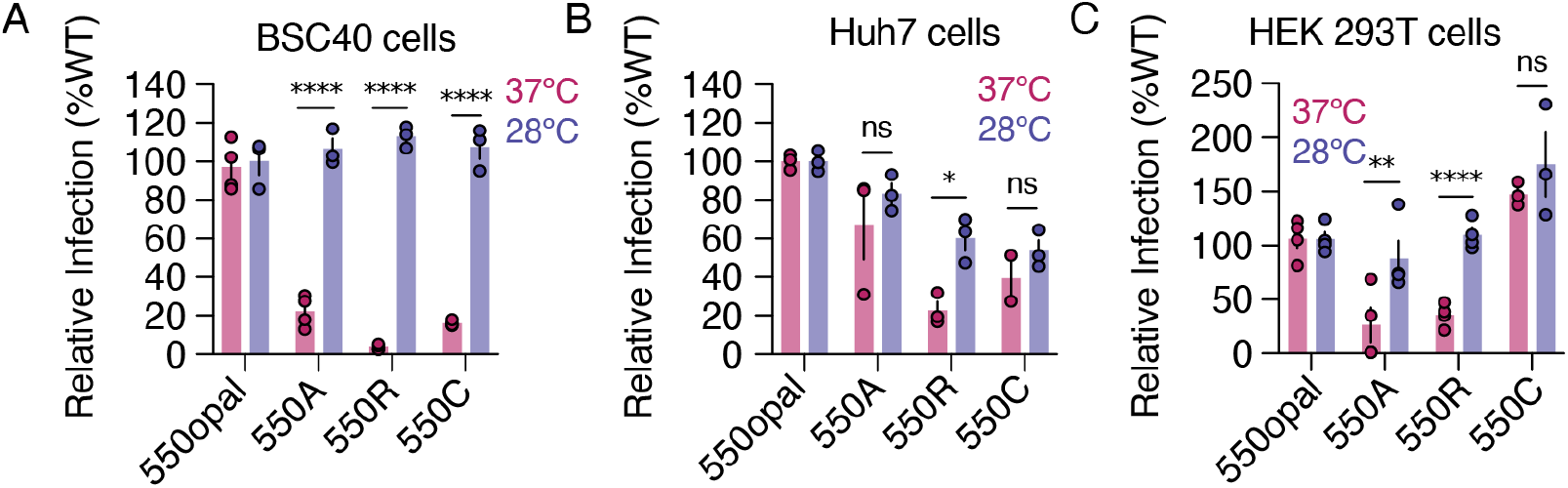
Infection rates of sense-codon SINV variants in different primate cells grown at either 37ºC or 28ºC. We transfected WT (550opal) or sense-codon (550A, 550R, or 550C) SINV variants expressing GFP into African green monkey (BSC40) cells (A), Human hepatocyte epithelial (Huh7) (B), or Human embryonic kidney (HEK 293T) cells (C) grown at either 37ºC or 28ºC. Cells were harvested 48 hours post-infection, and infection rates were quantified using flow cytometry. Two-way ANOVA with Tukey’s multiple comparisons test. ^****^ = P < 0.0001, ^**^ = P < 0.01, ^*^ = P < 0.05, ns = not significant.

**Suppl. Figure 6.**
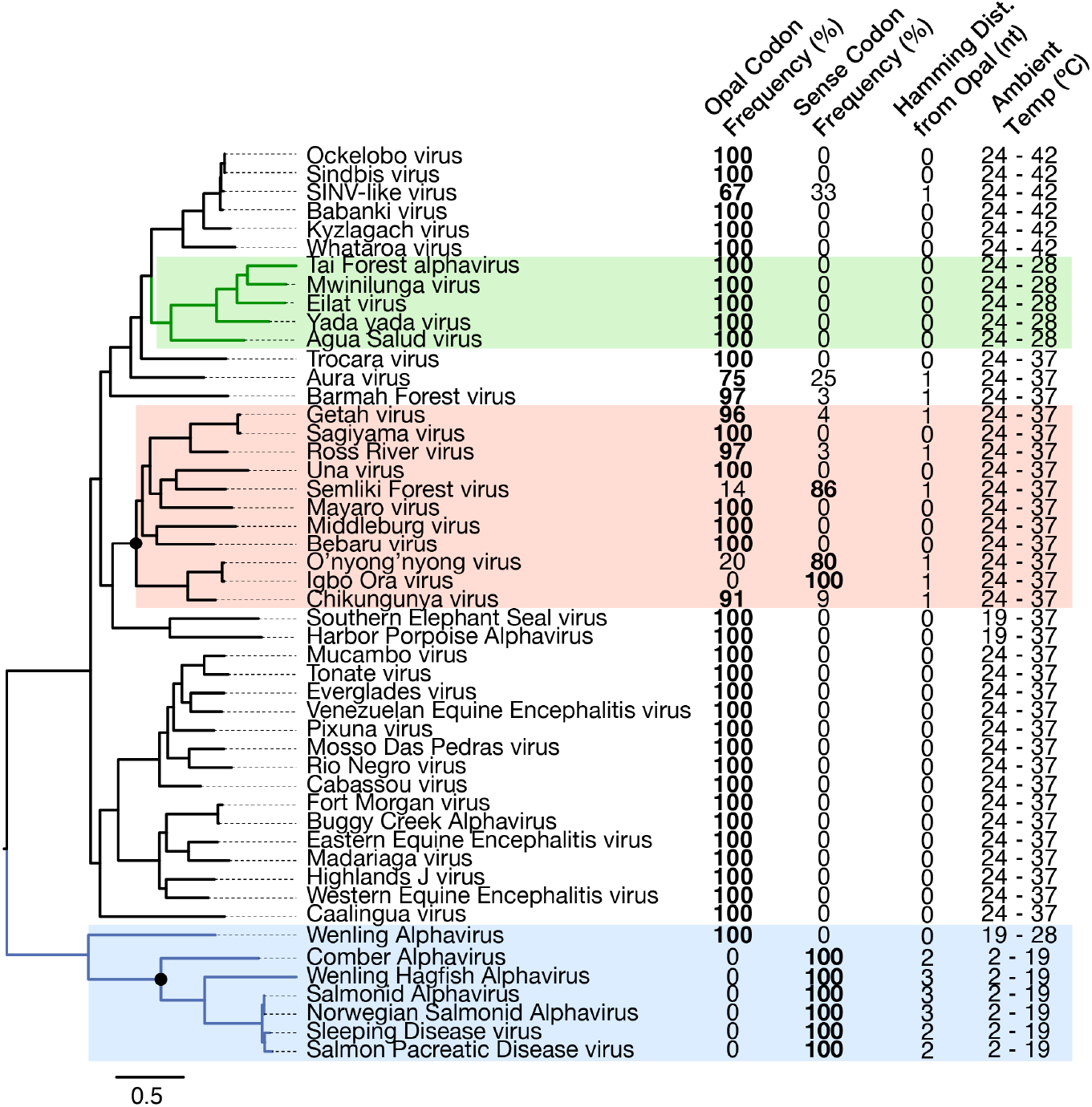
Phylogenetic conservation of the nsp3 opal codon in alphaviruses correlates with ambient temperature of hosts. A maximum likelihood phylogenetic tree of 49 extant alphaviruses. Dual-host viruses of the Semliki forest virus-clade are highlighted in light red. Single-host viruses like insect-restricted alphaviruses are highlighted in green and marine vertebrate-restricted alphaviruses are highlighted in blue. Nodes leading to lineages exhibiting a high incidence of sense-codons instead of opal are highlighted with solid black circles. Adjacent columns to each taxon indicate the percentage of sequenced isolates containing an opal stop codon (first column) versus a sense codon (second column) in the nsP3 gene. The third column shows the number of nucleotide differences between opal and sense codons. The fourth column shows the temperature ranges of the vertebrate or invertebrate hosts associated with each alphavirus species.

**Suppl. Table 1.**
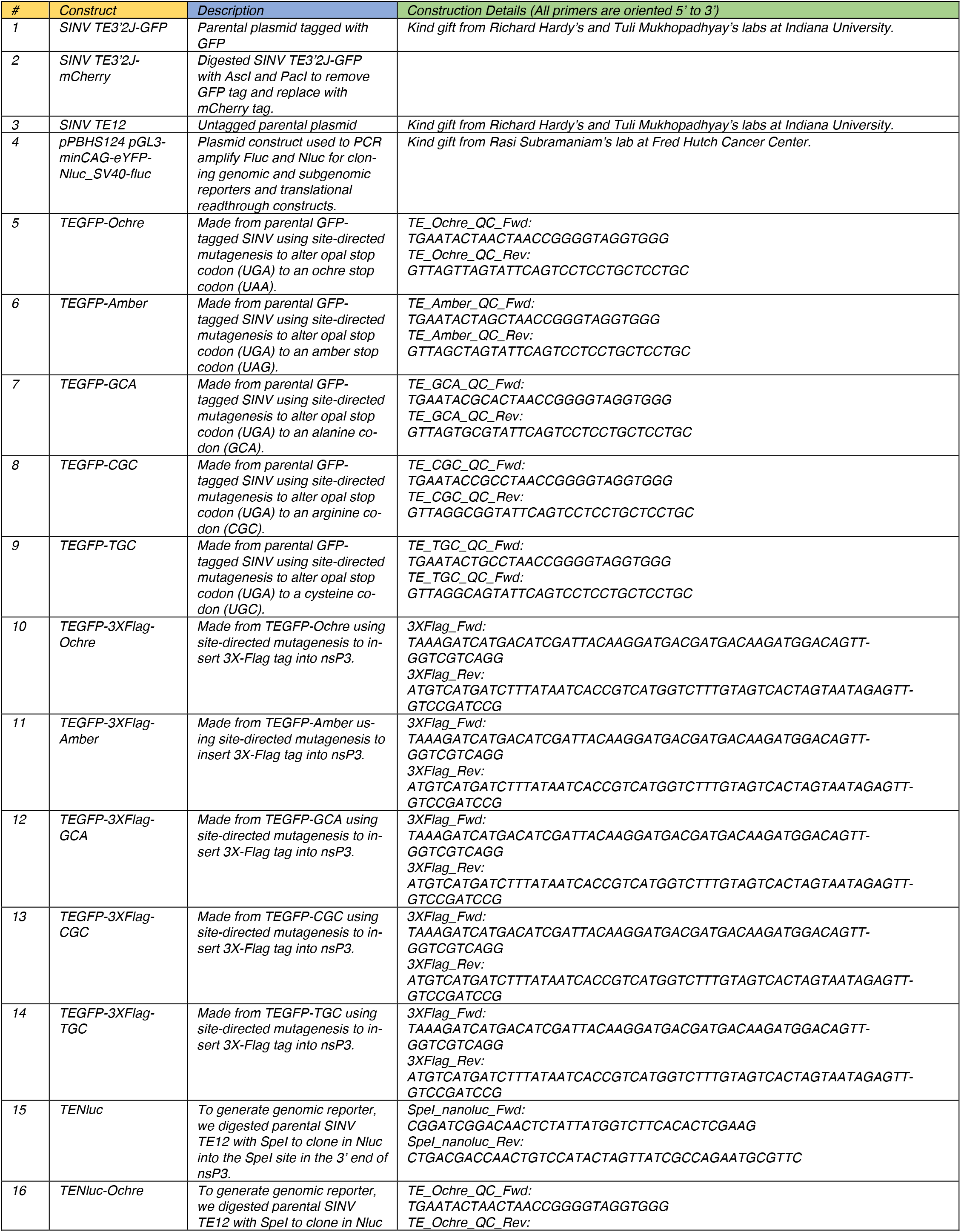
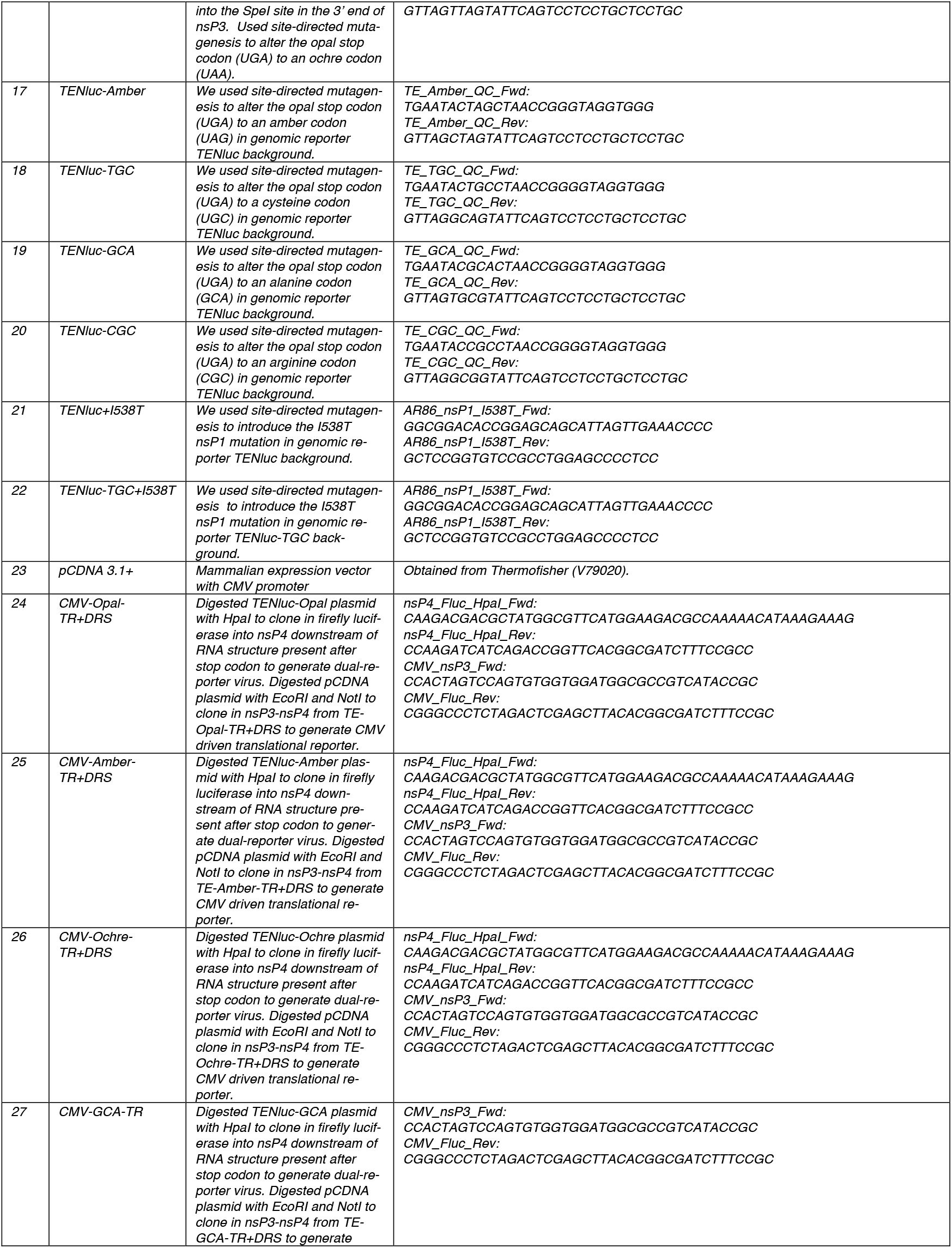
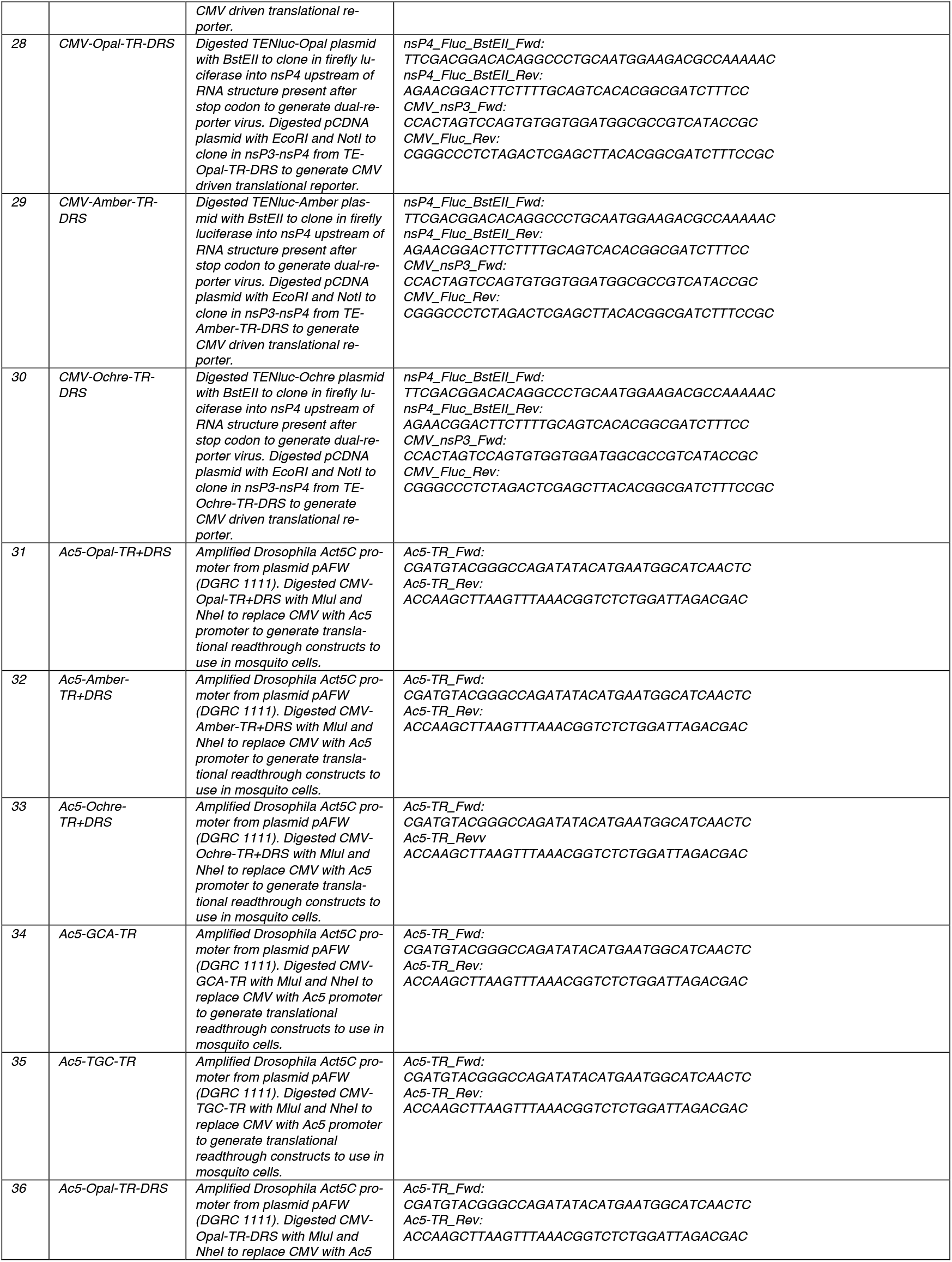
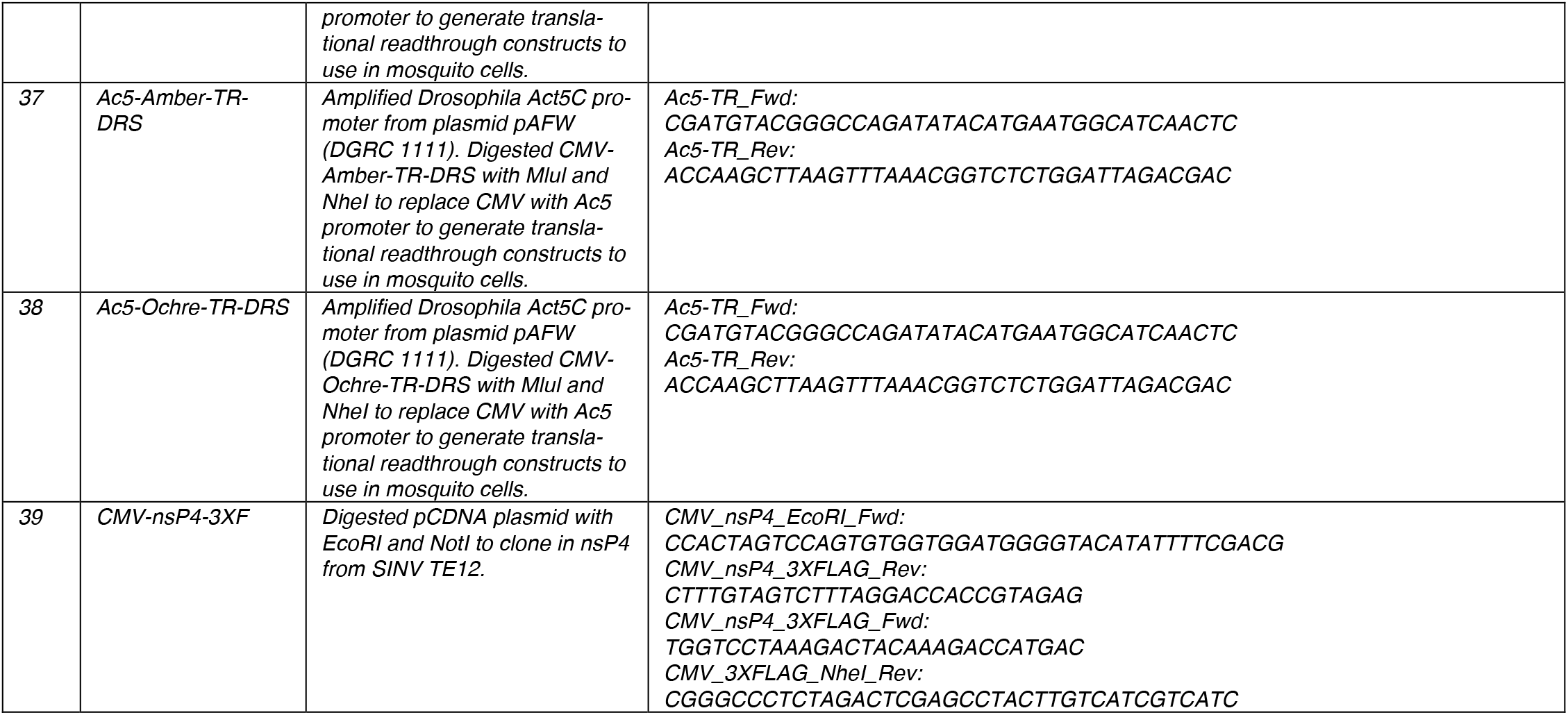
Constructs and primers used in this study.

## References

1. Levi, L. I. & Vignuzzi, M. Arthritogenic Alphaviruses: A Worldwide Emerging Threat? Microorganisms 7, 133 (2019).

2. Weaver, S. C., Winegar, R., Manger, I. D. & Forrester, N. L. Alphaviruses: Population genetics and determinants of emergence. Antiviral Res 94, 242–257 (2012).

3. Weaver, S. C. & Barrett, A. D. T. Transmission cycles, host range, evolution and emergence of arboviral disease. Nat Rev Microbiol 2, 789–801 (2004).

4. Ventoso, I. et al. Translational resistance of late alphavirus mRNA to eIF2α phosphorylation: a strategy to overcome the antiviral effect of protein kinase PKR. Genes Dev. 20, 87–100 (2006).

5. Morley, V. J. et al. Chikungunya virus evolution following a large 3’UTR deletion results in host-specific molecular changes in protein-coding regions. Virus Evol 4, vey012 (2018).

6. Strauss, E. G., Rice, C. M. & Strauss, J. H. Sequence coding for the alphavirus nonstructural proteins is interrupted by an opal termination codon. Proc. Natl. Acad. Sci. U.S.A. 80, 5271–5275 (1983).

7. Li, G. & Rice, C. M. The signal for translational readthrough of a UGA codon in Sindbis virus RNA involves a single cytidine residue immediately downstream of the termination codon. Journal of Virology 67, 5062–5067 (1993).

8. Palma, M. & Lejeune, F. Deciphering the molecular mechanism of stop codon readthrough. Biological Reviews 96, 310–329 (2021).

9. Strauss, E. G. & Strauss, J. H. Structure and Replication of the Alphavirus Genome. in The Togaviridae and Flaviviridae (eds. Schlesinger, S. & Schlesinger, M. J.) 35–90 (Springer New York, Boston, MA, 1986). doi:10.1007/978-1-4757-0785-4_3.

10. Shirako, Y. & Strauss, J. H. Regulation of Sindbis virus RNA replication: uncleaved P123 and nsP4 function in minus-strand RNA synthesis, whereas cleaved products from P123 are required for efficient plus-strand RNA synthesis. J Virol 68, 1874–1885 (1994).

11. Myles, K. M., Kelly, C. L. H., Ledermann, J. P. & Powers, A. M. Effects of an Opal Termination Codon Preceding the nsP4 Gene Sequence in the O’Nyong-Nyong Virus Genome on Anopheles gambiae Infectivity. J Virol 80, 4992–4997 (2006).

12. Jones, J. E. et al. Disruption of the Opal Stop Codon Attenuates Chikungunya Virus-Induced Arthritis and Pathology. mBio 8, 10.1128/mbio.01456-17 (2017).

13. Heise, M. T., Simpson, D. A. & Johnston, R. E. A Single Amino Acid Change in nsP1 Attenuates Neurovirulence of the Sindbis-Group Alphavirus S.A.AR86. Journal of Virology 74, 4207–4213 (2000).

14. Weaver, S. C., Brault, A. C., Kang, W. & Holland, J. J. Genetic and Fitness Changes Accompanying Adaptation of an Arbovirus to Vertebrate and Invertebrate Cells. Journal of Virology 73, 4316–4326 (1999).

15. Lanciotti, R. S. et al. Emergence of Epidemic O’nyongnyong Fever in Uganda after a 35-Year Absence: Genetic Characterization of the Virus. Virology 252, 258–268 (1998).

16. Mounce, B. C. et al. Chikungunya Virus Overcomes Polyamine Depletion by Mutation of nsP1 and the Opal Stop Codon To Confer Enhanced Replication and Fitness. Journal of Virology 91, 10.1128/jvi.00344-17 (2017).

17. Li, R., Sun, K., Tuplin, A. & Harris, M. A structural and functional analysis of opal stop codon translational readthrough during Chikungunya Virus replication. 2023.07.24.550286 Preprint at 10.1101/2023.07.24.550286 (2023).

18. Hwang Kim, K., Rümenapf, T., Strauss, E. G. & Strauss, J.H. Regulation of Semliki Forest virus RNA replication: a model for the control of alphavirus pathogenesis in invertebrate hosts. Virology 323, 153–163 (2004).

19. Rozen-Gagnon, K. et al. Alphavirus Mutator Variants Present Host-Specific Defects and Attenuation in Mammalian and Insect Models. PLOS Pathogens 10, e1003877 (2014).

20. Cereghino, C. et al. The E2 glycoprotein holds key residues for Mayaro virus adaptation to the urban Aedes aegypti mosquito. PLoS Pathog 19, e1010491 (2023).

21. Adelman, Z. N. et al. Cooler Temperatures Destabilize RNA Interference and Increase Susceptibility of Disease Vector Mosquitoes to Viral Infection. PLOS Neglected Tropical Diseases 7, e2239 (2013).

22. Lane, W. C. et al. The Efficacy of the Interferon Alpha/Beta Response versus Arboviruses Is Temperature Dependent. mBio 9, 10.1128/mbio.00535-18 (2018).

23. Athey, J. et al. A new and updated resource for codon usage tables. BMC Bioinformatics 18, 391 (2017).

24. Alexaki, A. et al. Codon and Codon-Pair Usage Tables (CoCoPUTs): Facilitating Genetic Variation Analyses and Recombinant Gene Design. J Mol Biol 431, 2434–2441 (2019).

25. Beier, H. & Grimm, M. Misreading of termination codons in eukaryotes by natural nonsense suppressor tRNAs. Nucleic Acids Research 29, 4767–4782 (2001).

26. Dabrowski, M., Bukowy-Bieryllo, Z. & Zietkiewicz, E. Translational readthrough potential of natural termination codons in eucaryotes – The impact of RNA sequence. RNA Biology 12, 950–958 (2015).

27. Kendra, J. A. et al. Functional and structural characterization of the chikungunya virus translational recoding signals. Journal of Biological Chemistry 293, 17536–17545 (2018).

28. Firth, A. E., Wills, N. M., Gesteland, R. F. & Atkins, J. F. Stimulation of stop codon readthrough: frequent presence of an extended 3′ RNA structural element. Nucleic Acids Res 39, 6679–6691 (2011).

29. Lemm, J. A., Durbin, R. K., Stollar, V. & Rice, C. M. Mutations which alter the level or structure of nsP4 can affect the efficiency of Sindbis virus replication in a host-dependent manner. Journal of Virology 64, 3001–3011 (1990).

30. Li, G. P. & Rice, C. M. Mutagenesis of the in-frame opal termination codon preceding nsP4 of Sindbis virus: studies of translational readthrough and its effect on virus replication. J Virol 63, 1326–1337 (1989).

31. de Groot, R. J., Rümenapf, T., Kuhn, R. J., Strauss, E. G. & Strauss, J. H. Sindbis virus RNA polymerase is degraded by the N-end rule pathway. Proceedings of the National Academy of Sciences 88, 8967–8971 (1991).

32. De Groot, R. J., Hardy, W. R., Shirako, Y. & Strauss, J. H. Cleavage-site preferences of Sindbis virus polyproteins containing the non-structural proteinase. Evidence for temporal regulation of polyprotein processing in vivo. The EMBO Journal 9, 2631–2638 (1990).

33. Rupp, J. C., Sokoloski, K. J., Gebhart, N. N. & Hardy, R. W. Alphavirus RNA synthesis and non-structural protein functions. Journal of General Virology 96, 2483–2500 (2015).

34. Lemm, J. A., Rümenapf, T., Strauss, E. G., Strauss, J. H. & Rice, C. M. Polypeptide requirements for assembly of functional Sindbis virus replication complexes: a model for the temporal regulation of minus- and plus-strand RNA synthesis. The EMBO Journal 13, 2925–2934 (1994).

35. Shin, G. et al. Structural and functional insights into alphavirus polyprotein processing and pathogenesis. Proc. Natl. Acad. Sci. U.S.A. 109, 16534–16539 (2012).

36. De Groot, R. J., Hardy, W. R., Shirako, Y. & Strauss, J. H. Cleavage-site preferences of Sindbis virus polyproteins containing the non-structural proteinase. Evidence for temporal regulation of polyprotein processing in vivo. The EMBO Journal 9, 2631–2638 (1990).

37. Carrasco, L., Sanz, M. A. & González-Almela, E. The Regulation of Translation in Alphavirus-Infected Cells. Viruses 10, 70 (2018).

38. Sokoloski, K. J. et al. Identification of Interactions between Sindbis Virus Capsid Protein and Cytoplasmic vRNA as Novel Virulence Determinants. PLOS Pathogens 13, e1006473 (2017).

39. Bhattacharya, T., Newton, I. L. & Hardy, R. W. Wolbachia elevates host methyltransferase expression to block an RNA virus early during infection. PLoS pathogens 13, e1006427 (2017).

40. Jose, J., Taylor, A. B. & Kuhn, R. J. Spatial and Temporal Analysis of Alphavirus Replication and Assembly in Mammalian and Mosquito Cells. mBio 8, 10.1128/mbio.02294-16 (2017).

41. King, C.-C. et al. Effect of incubation time on the generation of defective-interfering particles during undiluted serial passage of sindbis virus in Aedes albopictus and chick cells. Virology 96, 229–238 (1979).

42. Simpson, D. A., Davis, N. L., Lin, S.-C., Russell, D. & Johnston, R. E. Complete Nucleotide Sequence and Full-Length cDNA Clone of S.A.AR86, a South African Alphavirus Related to Sindbis. Virology 222, 464–469 (1996).

43. Baxter, V. K. & Heise, M. T. Immunopathogenesis of alphaviruses. in Advances in Virus Research vol. 107 315–382 (Elsevier, 2020).

44. Simmons, J. D., Wollish, A. C. & Heise, M. T. A Determinant of Sindbis Virus Neurovirulence Enables Efficient Disruption of Jak/STAT Signaling. J Virol 84, 11429–11439 (2010).

45. Heise, M. T. et al. An Attenuating Mutation in nsP1 of the Sindbis-Group Virus S.A.AR86 Accelerates Nonstructural Protein Processing and Up-Regulates Viral 26S RNA Synthesis. J Virol 77, 1149–1156 (2003).

46. Bartholomeeusen, K. et al. A Chikungunya Virus trans-Replicase System Reveals the Importance of Delayed Non-structural Polyprotein Processing for Efficient Replication Complex Formation in Mosquito Cells. J Virol 92, e00152–18 (2018).

47. Loh, E., Righetti, F., Eichner, H., Twittenhoff, C. & Narberhaus, F. RNA Thermometers in Bacterial Pathogens. Microbiology Spectrum 6, 10.1128/microbiolspec.rwr-0012–2017 (2018).

48. Meyer, A. et al. An RNA Thermometer Activity of the West Nile Virus Genomic 3′-Terminal Stem-Loop Element Modulates Viral Replication Efficiency during Host Switching. Viruses 12, 104 (2020).

49. Bisht, K. & te Velthuis, A. J. W. Decoding the Role of Temperature in RNA Virus Infections. mBio 13, e02021–22 (2022).

50. Hofmann, S., Cherkasova, V., Bankhead, P., Bukau, B. & Stoecklin, G. Translation suppression promotes stress granule formation and cell survival in response to cold shock. MBoC 23, 3786–3800 (2012).

51. Namy, O., Duchateau-Nguyen, G. & Rousset, J.-P. Translational readthrough of the PDE2 stop codon modulates cAMP levels in Saccharomyces cerevisiae. Molecular Microbiology 43, 641–652 (2002).

52. Forrester, N. L. et al. Genome-Scale Phylogeny of the Alphavirus Genus Suggests a Marine Origin. J Virol 86, 2729–2738 (2012).

53. Ryman, K. D. & Klimstra, W. B. Host responses to alphavirus infection. Immunological Reviews 225, 27–45 (2008).

54. Hickson, S. E. et al. Sequence diversity in the 3’ untranslated region of alphavirus modulates IFIT2-dependent restriction in a cell type-dependent manner. 2021.12.10.472177 Preprint at 10.1101/2021.12.10.472177 (2021).

55. Agranovsky, A. A. Where To Stop: Occurrence and Evolution of Translational Recoding Signals in RNA Viruses of Eukaryotes. Gene Expression 22, 240–249 (2023).

56. Bloom, J. D. Software for the analysis and visualization of deep mutational scanning data. BMC Bioinformatics 16, 168 (2015).

57. Wheeler, T. J., Clements, J. & Finn, R. D. Skylign: a tool for creating informative, interactive logos representing sequence alignments and profile hidden Markov models. BMC Bioinformatics 15, 7 (2014).

